# Spatially enforced engagement of PD-1⁺ T cells with AXL⁺ cells reprograms the tumor microenvironment and overcomes anti-PD-1 resistance

**DOI:** 10.64898/2026.10.26.761755

**Authors:** Ksenia M Yegodayev, Gregory Perelman, Tamar Shami, Bar Kaufman, Danialle Raviv, Divyasree Marripati, Aseel Khateeb, Raz Ben-uri, Oshrat Levi Galibov, Idan Milo, Alice Raizman, Alona Veinblat, Abraham Nissani, Noam Zuri, Dinesh Babu Manikandan, Ekaterina Rudnitsky, Ofra Novoplansky, Jonatan Zorea, Sarah Adi Eisdorfer, Sankar Jagadeeshan, Gili Ezri, Kamran Waidha, Raghda Abu Shareb, Monica Wainer, Uzi Hadad, Orit Itzhaki, Ronnie Shapira-Frommer, Yochai Wolf, Merav Cohen, Dexin Kong, Hai Wang, Angel Porgador, Leeat Keren, Amir Aharoni, Yaron Carmi, Ofir Cohen, Moshe Elkabets

## Abstract

**Background:** AXL overexpression promotes tumor-cell plasticity, local immune suppression, and resistance to PD-1 blockade. Conventional anti-AXL and anti-PD-1 antibodies combinations act independently and do not ensure physical proximity between PD-1⁺ T cells and AXL⁺ cells within the immunosuppressive tumor microenvironment.

**Methods:** To address this limitation, we developed a non-tandem, tetravalent IgG bispecific antibody designed to simultaneously engage PD-1 and AXL (BiTE-AXL/PD-1). Its antitumor activity was evaluated using in vitro functional assays and humanized and immunocompetent head and neck squamous cell carcinoma (HNSCC) models and compared with the combination of monospecific anti-AXL and anti-PD-1 antibodies. Spatial multiplex imaging was used to characterize treatment-associated changes in the tumor microenvironment of an anti-PD-1-resistant murine model.

**Results:** BiTE-AXL/PD-1 enhanced T-cell activation, promoted AXL-dependent tumor–T-cell conjugation, and increased tumor-cell killing in vitro, outperforming the combination of the two monospecific antibodies. In humanized and immunocompetent HNSCC models, BiTE treatment significantly delayed tumor growth compared with the antibody combination and was well tolerated, with no overt toxicity. Spatial multiplex imaging revealed a shift from a fibroblast-dominated microenvironment characterized by exhausted CD8⁺ T cells toward dendritic-cell- and CD8⁺ T-cell-enriched niches associated with antitumor immune activity.

**Conclusion:** BiTE-AXL/PD-1 promotes physical proximity between PD-1⁺ T cells and AXL⁺ cells, enhances antitumor immune responses, and remodels the immunosuppressive tumor microenvironment. These findings support PD-1/AXL bispecific targeting as a promising therapeutic strategy to overcome resistance to PD-1 blockade in HNSCC.

## Introduction

Immunotherapy has revolutionized cancer treatment, particularly immune checkpoint inhibitors like pembrolizumab and nivolumab which target the programmed cell death protein 1 (PD-1)/programmed death-ligand 1 (PD-L1) pathway[1–3]. These agents have shown clinical activity in multiple malignancies, including recurrent and metastatic head and neck squamous cell carcinoma (HNSCC)[4–7], by reinvigorating exhausted T cells and restoring anti-tumor immunity[8–10]. However, approximately 60-70% of HNSCC patients fail to respond to anti-PD-1 therapy, and acquired resistance frequently develops even among initial responders[5–7,11], underscoring the urgent need for strategies that enhance the efficacy and durability of PD-1 blockade. Mechanisms underlying PD-1 blockade resistance are multifactorial and include dysfunctional antigen presentation, upregulation of alternative immune checkpoint pathways, insufficient T-cell infiltration, and immunosuppressive stromal reprogramming, collectively limiting the ability of anti-PD-1 antibodies (Abs) to restore effective anti-tumor immunity[12–14].

Emerging evidence has identified AXL, a receptor tyrosine kinase, as a key mediator of immune suppression and limited response to PD-1 blockade[15–23]. AXL plays a multifaceted role in diverse cancers, including lung, breast, melanoma, and HNSCC[24,25]. In these tumors, its immunosuppressive effects are highly local and often depend on direct cell-cell contact between the AXL-expressing tumor or stromal cells and neighboring lymphocytes or antigen-presenting cells. AXL is expressed across multiple cell types within the tumor microenvironment (TME), including malignant cells[26,27], tumor-associated macrophages (TAMs)[28–30] and dendritic cells (DCs)[28,31], where it orchestrates a multifaceted immunosuppressive program. In tumor cells, AXL promotes epithelial-mesenchymal transition (EMT) and upregulates PD-L1 and PD-L2 while downregulating ICAM1 expression, impairing NK and T-cell-mediated cytotoxicity[21,26,32]. Within the stromal compartment, AXL profoundly shapes the TME by modulating the stromal and immune cell phenotype and function, generating an immunosuppressive niche that restricts T-cell-mediated tumor control[18]. In DCs, AXL signaling impairs antigen presentation and co-stimulation, causing defective CD8⁺ T-cell priming[31], and drives TAMs toward an immunosuppressive M2-like phenotype[29,33]. Critically, AXL directly induces PD-L1 and PD-L2 expression on tumor and myeloid cells[16,17,22,34–36], creating a mechanistic link between AXL activity and PD-1-mediated T-cell suppression. Accordingly, AXL inhibition reduces PD-L1 and enhances sensitivity to anti-PD-1 therapy preclinically[23,37–41], whereas high AXL expression correlates with primary and acquired resistance to PD-1 blockade in patients[22,42]. Combining AXL inhibition with anti-PD-1 blockade enhances anti-tumor activity in preclinical cancer models, and early clinical data suggest that dual targeting is feasible and may provide clinical benefit[43].

Despite this promise, anti-AXL and anti-PD-1 therapies circulate independently and bind their targets stochastically, without ensuring physical engagement between PD-1⁺ effector T cells and AXL-expressing cells; this lack of enforced proximity may limit synergy. Bispecific T-cell engagers (BiTEs), which physically bridge T cells to target cells through dual antigen recognition, can enforce spatial proximity and local immune synapse formation, concentrating immunomodulatory effects at sites of AXL-driven suppression. Most BiTE constructs are composed of two single-chain variable fragments (scFvs) and utilize CD3e as the T-cell binding domain, enabling broad polyclonal T-cell activation[44–46], however, CD3-based BiTEs are associated with short circulating half-lives and with significant systemic toxicity from non-specific T-cell activation and cytokine release syndrome[47,48]. In contrast, targeting PD-1 may be more selective by preferentially engaging tumor-infiltrating, antigen-experienced T cells that express PD-1 as a consequence of prior activation and chronic stimulation within the TME[49,50], potentially enhancing efficacy while reducing off-target toxicity.

Given the diversity of immunosuppressive AXL-expressing cells in the TME[26–31] and the correlation between AXL levels and acquired resistance to PD-1 blockade[22,23,37,42], we hypothesized that BiTE simultaneously targeting AXL and PD-1 would physically link PD-1⁺ T cells to AXL⁺ cells, relieve PD-1-mediated inhibition, and antagonize AXL-driven immune suppression in a spatially coordinated manner. By enforcing physical proximity between these cell populations, such BiTE could concentrate immune activation precisely at sites where AXL-mediated suppression is most active, potentially overcoming the spatial and functional barriers that limit the efficacy of conventional combination therapy.

In this study, we developed a tetravalent AXL/PD-1 BiTE that bridges PD-1⁺ CD8⁺ T cells with AXL⁺ tumor and stromal cells. We systematically optimized the BiTE format by evaluating multiple scFv configurations and selected a non-tandem tetravalent design based on superior functional activity in PD-1 reporter assays. We demonstrated that BiTE treatment strengthened anti-tumor immunity relative to combined anti-PD-1 and anti-AXL monotherapies. Specifically, BiTE binds PD-1 with high affinity and AXL with moderate affinity, promotes tumor-T cell conjugate formation in an AXL-dependent manner, and induces robust T-cell activation and tumor cell lysis in vitro, and tumor growth delay in two PBMCs humanized xenograft and two syngeneic HNSCC models. Using high-resolution multiplex immunofluorescence and spatial analysis, we revealed that BiTE reorganizes the TME by promoting co-infiltration of DCs and CD8⁺ T cells throughout the tumor, with peaks at its mid-core. We also showed that BiTE increases Tumor-DC and Tumor-DC-CD8 communities, shifting the TME from cancer associated fibroblasts (CAF)-dominated, immune-exhausted states to immune-active, DC-rich niches that support sustained T-cell activation and proliferation. Our findings establish spatially enforced dual engagement of AXL and PD-1 as an effective strategy to reprogram local immune interactions and augment anti-tumor T-cell function within the TME, with potential implications for overcoming checkpoint-blockade resistance in AXL-expressing tumors.

## Materials and Methods

### 1. Construction of Monospecific and BiTE Abs

Anti-AXL and anti-PD-1 scFv sequences (5′*V_L_*-linker-*V_H_*-linker, **Table S1**) were synthesized by IDT as double-stranded DNA fragments.

#### Human mono-AXL and mono-PD-1

scFv were cloned via Gibson assembly (New England Biolabs, E2621L) into a modified pFUSE-hIgG1e3-Fc2 plasmid encoding an engineered human IgG1 Fc domain with substitutions minimizing Ab-Dependent Cellular Cytotoxicity (ADCC) and complement-dependent cytotoxicity (CDC) (IgG2 residues at positions 233-236 and IgG4 residues at positions 327, 330, and 331: E233P, L234V, L235A, ΔG236, A327G, A330S, P331S). Plasmids were transformed into *E. coli*, selected on low-salt LB agar with 25 µg/mL zeocin (InvivoGen, ant-zn-05), and screened via colony PCR (ALLin™ Red Taq Mastermix, HighQu, PCM0205) using primers A&B (**Table S3**). Positive clones were verified by sequencing (Ben-Gurion University Genetics Unit) using Primers A-C. Candidates were validated by WB and ELISA.

#### Bispecific Abs

(BiTE-1: PD-1-AXL-Fc; BiTE-2: AXL-PD-1-Fc; BiTE-3: PD-1-Fc-AXL) were generated by PCR amplification of monospecific templates followed by Gibson assembly. For BiTE-1, mono-AXL plasmid was linearized using KAPA HiFi DNA polymerase (Roche, KK2101) with primers D& E; PD-1 scFv was amplified using KOD Hot Start DNA polymerase (Merck, 71086-3) with primers F& G (**Table S3**). For BiTE-2, mono-PD-1 was linearized using primers H& E; AXL scFv was amplified using primers I& J (**Table S3**). For BiTE-3, mono-PD-1 was linearized at the 3′ end of the Fc domain using primers K& L; the linker-AXL scFv sequence in BiTE-1 was amplified using KOD Hot Start polymerase. Plasmids were transformed into electrocompetent cells and validated by colony PCR and sequencing.

#### Murine variants

Mu-mono-PD-1 and Mu-BiTE were constructed analogously. Two anti-mouse PD-1 scFvs (**Table S1**) were synthesized by IDT. For Mu-mono-PD-1, anti-human AXL sequence was excised from pFUSE framework using KAPA HiFi polymerase with primers O&P (**Table S3**) and replaced with anti-mouse PD-1 scFvs via Gibson assembly. Validated clones were converted to Mu-BiTE by linearizing anti-murine PD-1 plasmids at the 3′ end of the Fc domain (primers K&L) using KOD Hot Start polymerase and inserting the linker-AXL sequence amplified from human BiTE-1. Clones were validated by colony PCR, sequencing, WB, and ELISA.

### 2. Cell Culture

HEK293F (ATCC) cells were cultured in FreeStyle™ 293 Expression Medium (Thermo Fisher, 12338018) in a 37°C orbital shaker (125-135 rpm, 8% CO_2_). HEK293T (ATCC) cells were maintained in FreeStyle™ medium at 37°C with 5% CO_2_. The IcAR-PD-1 reporter line, human PD-1 fused to zeta chain domain to drive an interleukin-2 (IL-2) reporter[51], and corresponding wild-type (WT) cells were generated in-house and maintained in RPMI 1640 medium supplemented with 1% L-glutamine (200 mM), penicillin (100 units/mL), streptomycin (100 µg/mL), and 10% fetal bovine serum (FBS). Human HNSCC cell lines SCC47 and FaDu were obtained from Dr. Reidar Grenman at the Department of Otorhinolaryngology-Head & Neck Surgery, Turku University, Finland, and ATCC, respectively. Murine mEERL cells were obtained from Dr. Paola D Vermeer at the Cancer Biology Research Center, Sanford Research (Sioux Falls, South Dakota). Human Breast cancer cell line MDA-MB231 was procured from the ATCC. The murine HNSCC cell line 4NQO-T was established in-house by exposing wild-type mice to drinking water containing 50 µg/mL 4-nitroquinoline-1-oxide (4NQO; Sigma, N8141)[52]. Cancer cell lines were cultured at 37°C with 5% CO_2_ in DMEM supplemented with 1% L-glutamine, penicillin (100 units/mL), streptomycin (100 µg/mL), and 10% FBS. Stable AXL-knockdown variants (shAXL and sgAXL) were generated as previously described[53]. All cell lines were routinely screened for mycoplasma and treated with De-Plasma (TOKU-E, D022) when required.

### 3. Ab Production

Cells were transiently transfected with 1 µg/mL of each pFUSE DNA plasmid encoding either mono-AXL or mono-PD-1 variants using GeneTranIII (Biomiga, GT2211-01) for small-scale expression. Bispecific variants were produced by transient transfection of HEK293T cells. Briefly, 5 million cells/plate were seeded in 10 mL DMEM 24 hours prior to transfection; medium was replaced with 5 mL FreeStyle™ 293 Medium before adding 10 µg of pFUSE DNA and 30 µg PEI (Polysciences, #23966) prepared in 500 µL DMEM (15-minute incubation at RT). Medium was replaced with fresh FreeStyle™ 293 Expression Medium 4 hours post-transfection. Conditioned supernatants were harvested at 24, 48, 72, and 96 hours post-transfection. Small-scale murine variants followed identical protocols. Bulk productions of human BiTE-3, mono-AXL, and mono-PD-1 utilized Chinese Hamster Ovary (CHOs) cells via TurboCHO™ platform (GenScript). Large-scale Mu-mono-PD-1 and Mu-BiTE utilized CHOs cells from Bionitron.

### 4. Western Blotting

Supernatants or cell lysates prepared in lysis buffer with phosphatase and protease inhibitors (Stratech, B15001-BIT, Millipore Sigma, P2714-1BTL) were quantified using Bradford assay (Bio-Rad, 5000006), boiled in SDS buffer for 5 minutes, resolved by 10% SDS-PAGE, and transferred to PVDF membranes using a Trans-Blot® Turbo™ Transfer Pack (Bio-Rad, 1704157). Membranes were blocked for 1h with 5% bovine serum albumin (BSA; AMRESCO, 0332-TAM) in TBS-T (Tris-buffered saline with 0.1% Tween 20). For **Ab expression in cell supernatants,** membranes were incubated with primary Abs diluted in blocking buffer for 1h at RT: anti-Human-Fc-HRP (Jackson Laboratories, 109-005-098, 1:2,500), anti-His-HRP (Sigma-Aldrich, A7058-1VL, 1:2,000), anti-total AXL (Cell Signaling Technology, 8661, 1:1,000). For **on-target effects of Abs on AXL receptor,** cancer cells were treated with the Abs at defined time intervals, lysed, and membranes were incubated overnight at 4°C with primary Abs against total AXL (Cell Signaling Technology, 8661, 1:1,000) or actin (Cell Signaling Technology, 4967, 1:1,000). The following day, membranes were incubated for 1 hour with HRP-conjugated anti-rabbit secondary Ab (Jackson Laboratories, 711-035-152, 1:10,000). Target protein levels were normalized to actin expression.

Signals were visualized using ECL reagents (Cyanagen, XLS3.0100/XLS071.0250) on an Azure c300 system and analyzed using ImageJ.

### 5. ELISA and Functional Artificial Reporter Assays

Assays were performed at room temperature with intermediate PBS-T (0.05% Tween 20) washes. **For human PD-1 binding**, streptavidin-coated plates (NEB, N7021S, 10 µg/mL in PBS) captured biotinylated human PD-1 (ACROBiosystems, PD1-H82E4, 50 ng/mL). **For AXL binding**, MaxiSorp plates (Thermo Fisher Scientific, 442404) were coated with human AXL recombinant protein (ACROBiosystems, AXL-H5226, 2 µg/mL in PBS). After blocking with 3% skimmed milk, serial Ab dilutions were added and detected with anti-human Fc-HRP (Jackson Laboratories, 109-005-098, 1:10,000 in PBS + 1% BSA). **For bridging ELISAs**, MaxiSorp plates coated with human AXL recombinant protein, blocked with 3% skimmed milk and incubated with serum containing BiTE, followed by biotinylated human PD-1 (50 ng/mL) and streptavidin-HRP detection (Jackson Laboratories, 016-030-084, 0.05 µg/mL). Development used TMB substrate (Surmodics, TMBW-1000-01), stopped with H_2_SO_4_, and read at 450 nm (Epoch spectrophotometer). **For IcAR binding assays**, 96-well plates (Corning Costar, 3596) were pre-coated overnight at 4°C with Ab supernatants or purified Abs (1-25 nM); positive controls: Keytruda/Nivolumab (Soroka Medical Center; negative controls: mono-AXL or human IgG4 isotype (BioLegend, 40370)). On the following day, IcAR-PD-1 cells (1 × 10^5^/well) were added and incubated for 24 hours. **For reporter blocking assays**, A549 target cells (0.25 × 10^5^/well) were stimulated for 24 hours with 1,000 IU/mL human recombinant IFN-*γ* and TNF-*α* (PeproTech). IcAR-PD-1 cells (1 × 10^5^/well) were then co-cultured with the targets for 20–24 hours in the presence of unpurified Ab supernatants. **For functional simultaneous binding assays**, FaDu tumor cells and IcAR-PD-1 cells (1 × 10^5^/well) were co-cultured with BiTE, Durvalumab (10 µg/mL, Soroka Hospital) and anti-PD-L2 (BioLegend, 345502, 1 µg/mL). Secreted murine IL-2 was quantified using a murine IL-2 commercial sandwich ELISA kit (BioLegend) as previously described[51]

### 6. Flow cytometry

**Cell surface binding assays** were performed at 4°C in FACS buffer (1% FBS, 1 mM EDTA, 0.05% sodium azide in PBS), incubated with unpurified or purified Abs for 1 hour. Bound Abs were detected by incubation with Alexa Fluor® 488 anti-human Fc (Jackson Laboratories, 709-545-098, 0.75 µg/mL) for 30 minutes. Background controls contained only the secondary Ab. **For AXL expression,** human FaDu or HEK293T cells were blocked for 30 minutes with FcR human blocking reagent (Miltenyi Biotec, 130-059-901, 1:1,000) and stained with BV786 anti-human AXL (BD Biosciences, 747857, 0.5 µg/mL). Murine 4NQO-T were blocked with TruStain fcX™ (anti-mouse CD16/32) Fc blocker (BioLegend, 101319) and stained with APC anti-mouse AXL (Thermo Fisher Scientific, 17-1084-82) or mouse IgG2b-APC isotype control (Miltenyi Biotec, 130-122-976).

#### Splenocyte Binding Assay

To assess binding under mitogenic activation, wild-type mouse splenocytes were harvested, filtered through a 70 µm strainer, stimulated with 1 µg/mL Concanavalin A (ConA; L7647, Sigma-Aldrich) for 48 hours and treated with Mu-mono-PD-1 or Mu-BiTE before detection with Alexa Fluor® 488 anti-human Fc.

#### T-cell Isolation & Proliferation Assay

Freshly isolated CD8⁺ T-cells were labeled with 5 µM CFSE (Invitrogen) for 15 minutes at 37°C, washed, and incubated for 1 hour to remove excess dye. Cells were co-cultured for 3 days at a 1:1 ratio with primary DCs pulsed with 5 µM recombinant gp100 (25-33) peptide under three conditions: (i) DC only, (ii) DC + 5 µg/mL Mu-BiTE, or (iii) DC + monospecific combination (5 µg/mL each), followed by flow cytometric tracking of CFSE dilution. Primary **DCs were isolated** from peripheral blood collected via cardiac perfusion with HBSS + 10 mM EDTA, fractionated on a Histopaque-1077 Hybri-Max gradient (Sigma-Aldrich), and centrifuged at 400 rcf for 15 minutes without brakes. Tumor tissues were digested in RPMI 1640 containing 2 mg/mL collagenase IV and 2,000 U/mL DNase I (Sigma-Aldrich) for 40 minutes at 37°C (200 rpm) and passed through a 70 µm strainer. Spleens and lymph nodes were homogenized in cold HBSS containing 2% FBS and 5 mM EDTA and passed through a 70 µm strainer. CD11b^+^ cells were isolated from tumor lysates, peripheral blood, and draining lymph nodes (DLN) using magnetic CD11b MicroBeads and columns (Miltenyi Biotec) and cultured for 4 days in DMEM containing 50 ng/mL GM-CSF and 10 ng/mL IL-4 (PeproTech) to generate MoDCs. Alternatively, cDCs were isolated from spleen and DLN homogenates using CD11c MicroBeads (Miltenyi Biotec) and allowed to adhere for 2 hours at 37°C before use.

Matching **primary T cells were harvested** from spleens and lymph nodes of euthanized B6.Cg-Thy1*^a^*/Cy Tg(TcraTcrb)8Rest/J mice processed through a 70 µm strainer, enriched via Histopaque-1077 gradient, and isolated using anti-CD8 magnetic nanobeads (MojoSort™, BioLegend).

All samples were stained with DAPI to exclude dead cells, acquired using a Beckman CytoFLEX flow cytometer, and analyzed using FlowJo software.

### 7. In Vitro Killing, Proliferation, and Interaction Assays

For **tumor cell killing assays,** GFP-expressing HNSCC target cells (1 × 10^4^/well) were co-cultured with human PBMCs (isolated via Ficoll [Sigma-Aldrich, GE17-1440-02]) at a 1:10 target-to-effector ratio in supplemented PBMC medium (RPMI 1640, 10% human serum, 1% sodium pyruvate, 1% L-glutamine, 1% MEM non-essential amino acids, 1% penicillin/streptomycin, 1% HEPES and 200 IU/mL recombinant human IL-2) with 0.5 nM Ab treatments. Live-cell imaging was performed for 96 hours on a JuLI-stage platform (NanoEnTek) and analyzed using JuLI STAT software. Secreted IFN-*γ* was measured using a Human IFN-*γ* Standard TMB ELISA Development Kit (PeproTech, 900-T27) read at 450 nm and 620 nm using an Epoch spectrophotometer.

**Surface Plasmon Resonance (SPR)** affinities were resolved using a ProteOn™ XPR36 System (Bio-Rad). Purified Abs were immobilized via EDC/s-NHS chemistry on SPP CMD700L sensor prisms (XanTec bioanalytics) in acetate buffer to achieve response units (RU) of 596 (mono-AXL), 2,036 (mono-PD-1), and 1,645 (BiTE). Unreacted sites were deactivated with ethanolamine, followed by a PBST wash. Recombinant human AXL and PD-1 protein analytes (ACROBiosystems) were injected in PBST at varying concentrations. Sensors were regenerated between cycles using NaOH. Kinetic constants (*k_d_*, *k_a_*, and *K_D_*) were derived using a Langmuir binding model with double background subtraction

**Cellular bridging** was evaluated by imaging cell doublets on an ImageStream MKII platform (Cytek Amnis). FaDu-GFP cells and DiD-labeled IcAR-PD-1 cells were co-incubated at 4°C for 2 hours with 5 nM Ab treatments. Images were acquired using a 40X objective and analyzed via IDEAS® 6.2 software using an Area vs. Aspect Ratio brightfield gating strategy.

To verify **AXL-dependence in conjugate formation,** CFSE-labeled FaDu cells (WT or shAXL; BioLegend, 423801) were co-incubated with mCherry-labeled IcAR-PD-1 cells for 2 hours at 4°C with BiTE or monospecific combination and double-positive (CFSE^+^mCherry^+^) percentage within the live doublet population was determined by flow cytometry and FlowJo software.

For **antigen-specific T-cell killing**, CD8^+^ gp100-reactive T cells expanded for 3 days on 1 µM gp100-pulsed splenocytes isolated using Ficoll and anti-CD8 magnetic nanobeads (MojoSort™, BioLegend). Labeled T cells were co-cultured with gp100-pulsed tumor targets (1:4 ratio) for 16 hours ± 5 µg/mL of monospecific or Mu-BiTE Abs. Target cell apoptosis was determined by quantifying Caspase 3/7 activity via flow cytometry.

### 8. Animal models

Animal studies were approved by the Institutional Animal Care and Use Committee (IACUC) of Ben-Gurion University of the Negev (protocols IL-44-06-2019E and IL-64-10-2023E). Mice were housed in air-filtered laminar flow cabinets with a 12-hour light/dark cycle with ad libitum access to food and water. At the end of the experiments, animals were euthanized using CO_2_. NOD.Cg-PrkdcscidIl2rgtm1Wjl/SzJ (Nod scid gamma-NSG) mice were obtained from Jackson Laboratory (Bar Harbor, Maine, USA) and C57BL/6JRccHsd (WT) mice were purchased from Envigo (Indianapolis, Indiana, USA). All experiments utilized mice aged 6-10 weeks.

### 9. Human samples

Patients with metastatic melanoma who failed on anti-PD1 were recruited by the Ella Lemelbaum Institute for immuno-oncology and melanoma according to approval no. SMC-3566-16 approved by the Sheba Medical Center Ethics committee.

### 10. Pharmacokinetics, Biodistribution, and In Vivo Safety

#### Pharmacokinetics

BiTE was incubated in human serum (Type AB, Capricorn, HUM-3B) in a final concentration of 1 μM at 37°C for short-term (6 days) and long-term (1 month) or administered as a single i.p. injection (10 mg/kg) to naive NSG mice (*n* = 4). Samples were collected over a time course (for in vivo - via the tail vein using Microvette® CB 300 Z blood collection tubes (Sarstedt AG)) and processed via bridging ELISA and half-life parameters were extracted using a two-compartment pharmacokinetic model.

#### Biodistribution

Subcutaneous FaDu CDX tumors (2 × 10^6^cells in 100 µL PBS) were established in female NSG mice. Mice were randomized by tumor volume and injected i.p. with 5 mg/kg Alexa Fluor™ 680-labeled BiTE (*n* = 3) using a Saivi™ Rapid Ab Labeling Kit (Thermo Fisher Scientific, S30045) or PBS (*n* = 2). Daily near-infrared whole-body imaging was executed for 4 days under isoflurane anesthesia using a Newton 7.0 FT500 system (Vilber), and fluorescence signals were normalized ex vivo.

#### In Vivo Safety

Naive male NSG mice were engrafted i.v. with human PBMCs (3 × 10^6^ cells in 200 µL PBS) and treated i.p. twice weekly with vehicle (*n* = 3) or 10 mg/kg BiTE (*n* = 4) for 7 days. Post-euthanasia serum was analyzed for ALT, ALK-phos, blood urea, and creatinine at the Biochemical Laboratory of Soroka Medical Center.

### 11. In Vivo Therapeutic Efficacy

**Subcutaneous CDX model,** male NSG mice were injected s.c. with SCC47 cells (5 × 10^6^ cells in 100 µL PBS) into the flanks. One-week post-injection, mice were engrafted i.v. with healthy donor human PBMCs (9 × 10^6^ cells in 200 µL PBS) and randomized into three treatment arms (*n* = 7–8 mice per group): monospecific combination (mono-AXL + mono-PD-1, 2 mg/kg each), BiTE (2 mg/kg), or PBS vehicle. Treatments were administered i.p. twice weekly. A second i.v. injection of PBMCs was administered one week later.

**Orthotopic CDX model**, male NSG mice were injected s.c. with FaDu cells (2 × 10^6^ cells in 50 µL PBS) into the lip, followed one week later by i.v. PBMC engraftment (1 × 10^7^ cells) and twice-weekly i.p. treatments of monospecific combination (5 mg/kg each) or molar-equivalent BiTE (7.6 mg/kg).

**Syngeneic dose efficacy assays:** male WT C57BL/6 mice bearing bilateral subcutaneous 4NQO-T tumors (5 × 10^6^ cells in 100 µL PBS) were randomized to receive twice-weekly i.p. dose-escalation injections of Mu-BiTE (0.1, 1, or 5 mg/kg) or PBS.

**Syngeneic comparative efficacy**, orthotopic lip models were established in male WT mice using either 2 × 10^6^ mEERL cells or 5 × 10^6^ 4NQO-T cells. Mice were randomized into three groups (*n* = 5 per group) and treated i.p. twice weekly. In the mEERL model, mice received either 5 mg/kg of each monospecific component or 5 mg/kg of Mu-BiTE. In the 4NQO-T model, treatments consisted of the monospecific combination (5 mg/kg each) or molar-equivalent Mu-BiTE (7.6 mg/kg).

Tumor volumes were calculated as: volume = length × width^2^ × (*π*/6).

### 12. Immunohistochemistry

Tissues were fixed in 4% paraformaldehyde (PFA) solution at room temperature for no more than 24 hours, dehydrated, and embedded in paraffin. Tissue sections were deparaffinized using xylene, and endogenous peroxidase activity was blocked by treating with 3% hydrogen peroxide (H₂O₂) for 20 minutes. Subsequently, sections were rinsed with water for 5 minutes. Antigen retrieval was carried out in citrate buffer (pH 6) at 99.9 °C for 15 minutes. The sections were then blocked for 1 hour at room temperature with a blocking solution composed of phosphate-buffered saline (PBS), 0.1% Tween, and 5% bovine serum albumin (BSA). Following blocking, the sections were incubated overnight at 4 °C with the primary antibody CD8 (CST-85336S, 1:200) and CD45 (CST-13917S, 1:200) diluted in blocking solution. On the next day, the slides were washed three times with PBS-Tween (PBS-T), and color detection was performed using an ABC kit (VECTASTAIN, Cat. VE-PK-6200) as per the manufacturer’s instructions. Sections were counterstained with hematoxylin and mounted using Micromount mounting medium (Leica, Cat. 380-1730). All slides were digitized using a Panoramic Scanner (3DHISTECH, Budapest, Hungary), and image analysis was performed with Qupath-0.2.3 software. Tumor tissue fields for analysis were selected by a blinded investigator. Cells in the selected fields were detected using Qupath-0.2.3 software and classified as positive or negative for DAB staining based on thresholds determined independently by two blinded investigators. Cell detection criteria and threshold values were standardized across all slides. To confirm the specificity of the staining and analysis thresholds, a matched negative control was used. This control was processed identically but without incubation with the primary antibody, ensuring all secondary antibody development procedures were followed.

### 13. Multiplex Spatial Imaging and Image Analysis

FFPE tissue sections (5 µm) were deparaffinized, rehydrated, and subjected to multiplex immunofluorescence using the Akoya Biosciences CODEX® platform[54]. Sections were stained with a customized panel of primary Abs conjugated to unique oligonucleotide barcodes (**Table S4**). Automated cyclical imaging was executed on a PhenoCycler-Fusion system (Akoya) using an Olympus 20X/0.7 objective.

**Images were analyzed** in QuPath (0.5.9[55]). Low-quality or artifact-heavy sections were annotated and excluded. Cell segmentation was performed using a StarDist[56] nuclear detection model on the DAPI channel and whole-cell masks were generated by isotropically expanding nuclear boundaries by 5 µm, bounded by adjacent cells. CD31-positive vasculature was reconstructed using spatial tessellates rather than individually segmented endothelial cells[57] because the elongated and interconnected morphology of endothelial cells was not reliably captured by the nucleus-centered single-cell segmentation used for other cell populations. For each sample, the CD31 channel was Gaussian-smoothed (σ = 1 pixel) and thresholded using manually curated thresholds were applied to generate a binary CD31-positive mask. The mask was divided into non-overlapping 12 × 12 µm tiles, and tiles containing ≥20% CD31-positive pixels were retained. The centroid of each retained tile was registered as a vessel-derived spatial proxy and incorporated into the cell-level metadata as an “Endotheliocyte” for downstream spatial analyses. These objects therefore represent local CD31-positive vascular regions rather than individually segmented endothelial cells.

**Single-cell phenotyping** was performed using mean fluorescence intensities extracted for all available markers. Phenotypic lineages were assigned by manual gating based on canonical lineage markers (Table S2). Cells lacking leukocyte or stromal marker expression were classified as putative malignant cells.

**Spatial zoning** was computed using custom Python scripts. A tissue mask established the structural centroid and defined a normalized radial depth axis from the border to the inner core, partitioning each sample into four 25% radial bands: Margin, Outer Core, Mid Core, and Deep Core. Zone-specific counts tracked cell localization gradients. Treatment-associated localization patterns were normalized using a **weighted spatial distribution metric**: P(s,g,c,z) = N(s,g,c,z) / Σ_z N(s,g,c,z) × 100, where N(s,g,c,z) is the number of cells of type c in zone z from sample s in treatment group g; thus, for each sample and cell type, the distribution across all zones summed to 100%. To correct for total abundance variation between treatment arms, frequencies were adjusted by global cellular prevalence within that treatment group: w(g,c) = N(g,c) / N(g), where N(g,c) is the total number of cells of type c across all samples in treatment group g and N(g) is the total number of cells in that group. The weighted zone distribution was then calculated as WP(s,g,c,z) = P(s,g,c,z) × w(g,c).

**Cellular communities** were identified by local cell-type composition profiles surrounding individual cells[58], defined as the fractions of neighboring cells belonging to each cell type within a 40 µm radius of each cell centroid. These neighborhood composition vectors were computed using the scimap[59] package. Neighborhood vectors were then clustered using MiniBatchKMeans (scikit-learn package)[60], iterating the number of clusters *k* from 10 to 20 to select the solution maximizing the cell-level silhouette score.

**Relative community abundance** was computed across treatment arms as follows: for each community c, the relative contribution of each treatment group g was calculated as R(g,c) = N(g,c) / Σ_g N(g,c) × 100, where N(g,c) is the number of cells from treatment group g assigned to community c; thus, the contributions of all treatment groups summed to 100% within each community. The communities were ranked according to their relative representation in BiTE-treated tumors.

**Functional heatmaps** (activation/exhaustion), marker-positive percentages were calculated out of the total CD8^+^ T cell population per designated zone or community.

### 14. Statistical Analysis

Data were evaluated using GraphPad Prism (v9) or Python, and values are presented as mean ± SEM or mean ± SD. Group differences were evaluated via Student’s t-tests or one-way/two-way ANOVA followed by Tukey’s or Sidak’s post hoc tests. Continuous in vivo tumor growth curves were statistically compared using permutation tests (http://bioinf.wehi.edu.au/software/compareCurves/index.html).

For multiplex spatial mapping, localized features were quantified at the level of local tissue tiles while retaining the biological specimen (*n* = 3) as the unit of replication. Each specimen was partitioned into a 5 × 5 grid of equal-area tiles (tiles with < 50 cells were excluded). Compositional differences between treatment groups were tested using binomial generalized linear models with a logit link, modeling specific counts out of the total cell index per tile. To mitigate spatial pseudo-replication, standard errors were clustered at the biological sample level.

Data are labeled with exact p-values in individual plots. In the case of dense multiple-comparison graphs, only significant differences are shown, and non-significant comparisons (p ≥ 0.05) are omitted.

## Results

### Development and characterization of a non-tandem tetravalent bispecific antibody targeting AXL and PD-1

To enhance anti-tumor immunity via spatially directed dual-targeting, we developed a BiTE antibody (Ab) capable of engaging AXL on tumor cells and PD-1 on CD8⁺ T cells. First, we generated single-arm monospecific Abs targeting AXL or PD-1 individually (**Fig. 1A**). Abs were constructed with single-chain variable fragments (scFv) by linking the variable regions of the heavy and light chains through a flexible peptide linker, selecting candidate scFv sequences from previously reported anti-AXL and anti-PD-1 Abs (**Table S1**). We fused each scFv to a mutated Fc domain to prolong the half-life while eliminating ADCC generating two scFv variants per target (**Fig. S1A**). Based on binding capacity evaluated using enzyme-linked immunosorbent assay (ELISA) (**Fig. S1B**) and flow cytometry (**Fig. S1C**), we selected the optimal scFv variants for targeting AXL and PD-1 separately (AXL scFv2 and PD-1 scFv2, respectively), hereafter referred to as mono-AXL and mono-PD-1, respectively. Second, we designed three tetravalent Quadra-BiTE configurations by fusing tandem scFv and mutated Fc backbone to test how spatial arrangement influences performance: two tandem scFv BiTEs (BiTE-1, BiTE-2) and one non-tandem BiTE with scFvs on both sides of the Fc domain (BiTE-3) (**Fig. S1D**). These formats were assessed for retention of binding capacity relative to the corresponding mono-Abs using ELISA (**Fig. S1E**) and flow cytometry **(Fig. S1F)**. To functionally assess the anti-PD-1 arm, we used the engineered PD-1 reporter T-cell system IcAR-PD-1[51], which quantifies receptor activation through interleukin-2 (IL-2) secretion. BiTE-3 displayed strongest PD-1-dependent activation despite the lowest binding capacity of the three formats (**Fig. 1B**). Therefore, we selected this non-tandem tetravalent configuration (BiTE-3), hereafter referred to as BiTE **(Fig. 1A**). Third, we characterized BiTE and its corresponding mono control Abs following large-scale production. Reduced (R) and non-reduced (NR) SDS-PAGE revealed monomers of ∼60 kDa for the mono-Abs and ∼75 kDa for BiTE, and dimers of ∼120 kDa and ∼150 kDa, respectively (**Fig. S1G**), with 95-97% purity by size-exclusion high-performance liquid chromatography (SEC-HPLC) (**Fig. S1H**). To assess binding kinetics and affinities via surface plasmon resonance (SPR), we immobilized the Abs on a prism and introduced recombinant AXL and PD-1 proteins at various concentrations. Mono-AXL bound AXL, with a *K_D_* of 20 nM, whereas BiTE bound with a *K_D_* of 60 nM. For PD-1, mono-PD-1 and BiTE were comparable, with a *K_D_* of approximately 6 nM **(Fig. 1C**). BiTE showed slow dissociation from PD-1 (*K_d_* ∼ 7 × 10^−4^ 1/sec) suggesting stable binding, consistent with IcAR-PD-1 activation. Flow cytometry analysis of binding to AXL receptor on FaDu cancer cells demonstrated dose-dependent binding for both Abs with higher apparent binding potency observed for mono-AXL compared with BiTE, consistent with ELISA results (**Fig. 1D**). In AXL-knockdown FaDu cells **(Fig. S1I)**, both Abs exhibited reduced binding (**Fig. 1E**) confirming AXL specificity. As a test of the AXL arm, we examined the effect of Ab treatment on AXL protein levels as previously described[61]. Both mono-AXL and BiTE induced a time-dependent reduction in total AXL in two AXL-high cell lines, FaDu and MDA-MB-231 (**Fig. 1F**). For the PD-1 arm, mono-PD-1 and BiTE bound IcAR-PD-1 cells in a dose-dependent manner (**Fig. 1G**), with negligible binding to low PD-1 IcAR-WT cells (**Fig. 1H)**. IL-2 secretion showed that both mono-PD-1 and BiTE were as effective as the clinically approved anti-PD-1 antibody nivolumab in stimulating the PD-1 receptor (**Fig. 1I**). Together, these analyses establish BiTE as a well-assembled, target-specific bispecific Ab that retains AXL engagement, PD-1 blockade, and T-cell activation capacity.

**Figure 1.**
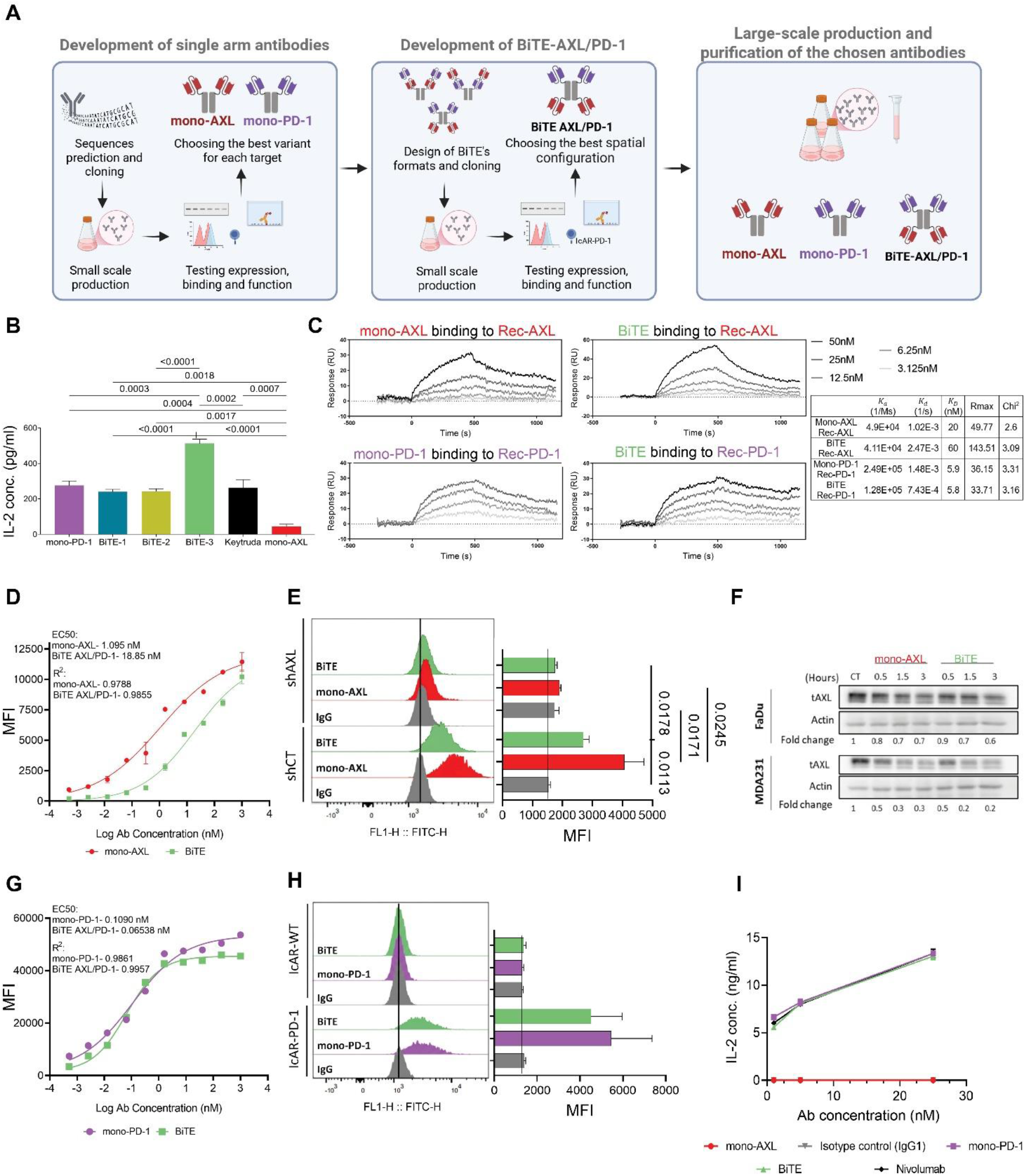
Development and characterization of a non-tandem tetravalent bispecific antibody targeting AXL and PD-1. **(A)** Development workflow of BiTE-AXL/PD-1 antibody. **(B)** IcAR-PD-1 binding assay of anti-PD-1 (IL-2 concentration measured by ELISA) for unpurified mono-PD-1, BiTE variants and mono-AXL (negative control); Keytruda (10 µg/mL) as positive control. Mean ± SEM of technical triplicates (n = 3) from single experiment. Statistical significance was determined using a one-way ANOVA followed by two-tailed Tukey’s multiple comparisons test with correction for multiple comparisons (adjusted P values shown). **(C)** SPR results showing binding of recombinant AXL to mono-AXL and BiTE antibodies, and binding of recombinant PD-1 to mono-PD-1 and BiTE antibodies. Grey shades represent the recombinant protein concentrations (3.125 nM, 6.25 nM, 12.5 nM, 25 nM and 50 nM). Single experiment. **(D)** Dose-response binding of mono-AXL and BiTE to FaDu cells by flow cytometry. Background-subtracted gMFI versus log10 concentration, EC50 calculated from a four-parameter fit. Representative of two biological experiments; mean ± SEM of technical duplicates (n = 2). **(E)** Abs binding to FaDu shCT and shAXL cells by flow cytometry. Log scale (FL1), MFI at right; representative of two biological experiments; mean ± SEM of biological duplicates (n=2). Statistical significance was determined using a one-way ANOVA followed by two-tailed Tukey’s multiple comparisons test (adjusted P values shown). **(F)** WB of total AXL (tAXL) over time in FaDu and MDA-MB-231 cells treated with control, mono-AXL, or BiTE. Protein levels were quantified by densitometry and normalized to actin. Fold-change values are shown relative to the corresponding control time points. Representative of two biological experiments (n = 2). **(G)** Same as D for mono-PD-1 and BiTE to IcAR-PD-1 cells. **(H)** Same as E to IcAR-PD-1 (PD-1-high) and IcAR-WT (PD-1-low) cells. **(I)** IcAR-PD-1 functional assay: IL-2 secretion measured for plates coated with various concentrations (1 nM, 5 nM and 25 nM) with mono-PD-1 or BiTE; isotype control and mono-AXL as negative controls, nivolumab as positive control; representative of two biological experiments; mean ± SEM of technical duplicates (n = 2).

### BiTE promotes tumor-T-cell interaction and enhances T-cell activation and cytotoxicity in an AXL-dependent manner

To test whether BiTE enhances physical interaction between T cells and AXL-expressing cells, we performed a co-culture bridging assay. GFP-labeled AXL-expressing FaDu cells, were co-incubated with DiD-labeled IcAR-PD-1 cells and treated with BiTE, a combination of mono-AXL and mono-PD-1 (Combo), or BiTE with competing mono-PD-1 and mono-AXL (**Fig. 2A)**. Imaging flow cytometry showed that BiTE significantly increased tumor-T-cell doublets with the GFP-DiD tumor-T-cell population showing more than a two-fold increase following BiTE treatment (**Fig. 2B** and **S2A**). Competition with mono-AXL or mono-PD-1 markedly reduced doublet formation, indicating target-dependent bridging (**Fig. 2B**). Image analysis confirmed BiTE-mediated tumor-immune interactions (**Fig. 2C**). Bridging experiments using AXL-knockdown cells (FaDu shAXL) (**Fig. S1I**) showed fewer doublets with FaDu shAXL cells than with the FaDu shCT cells, indicating that both arms contribute to cancer-immune doublet formation (**Fig. 2D**). To assess sustained anti-PD1 activity during bridging, we measured IL-2 in the IcAR-PD-1 system, with and without PD-L1/PD-L2 blockade (**Fig. 2E**). BiTE increased IL-2 levels relative to untreated co-cultures, and this enhancement persisted under PD-L1/PD-L2 blockade (**Fig. 2F**), demonstrating PD-1-dependent T-cell activation. We next assessed T-cell engagement and tumor killing in several co-culture systems. We co-cultured peripheral blood mononuclear cells (PBMCs) with GFP-labeled SCC47 cells in the presence of BiTE or Combo (**Fig. 2G**). BiTE significantly increased tumor cell killing compared to controls and induced significantly higher IFN-γ secretion than vehicle with also a trend of higher IFN-γ secretion compared to Combo (**Fig. 2G**). Similar enhanced cytotoxic activity was also observed in FaDu co-culture (**Fig. S2B**). In a patient-derived system, tumor-infiltrating lymphocytes (TILs) co-cultured with AXL-expressing melanoma cells (**Fig. S2C**) produced significantly more IFN-γ with BiTE than Combo (**Fig. 2H**) whereas low-AXL melanoma cells showed no such enhancement (**Fig. S2D**), supporting AXL-dependent T-cell activation. Altogether, the findings show that BiTE drives AXL-dependent tumor-T-cell conjugation and promotes T-cell activation and tumor killing across multiple systems.

**Figure 2.**
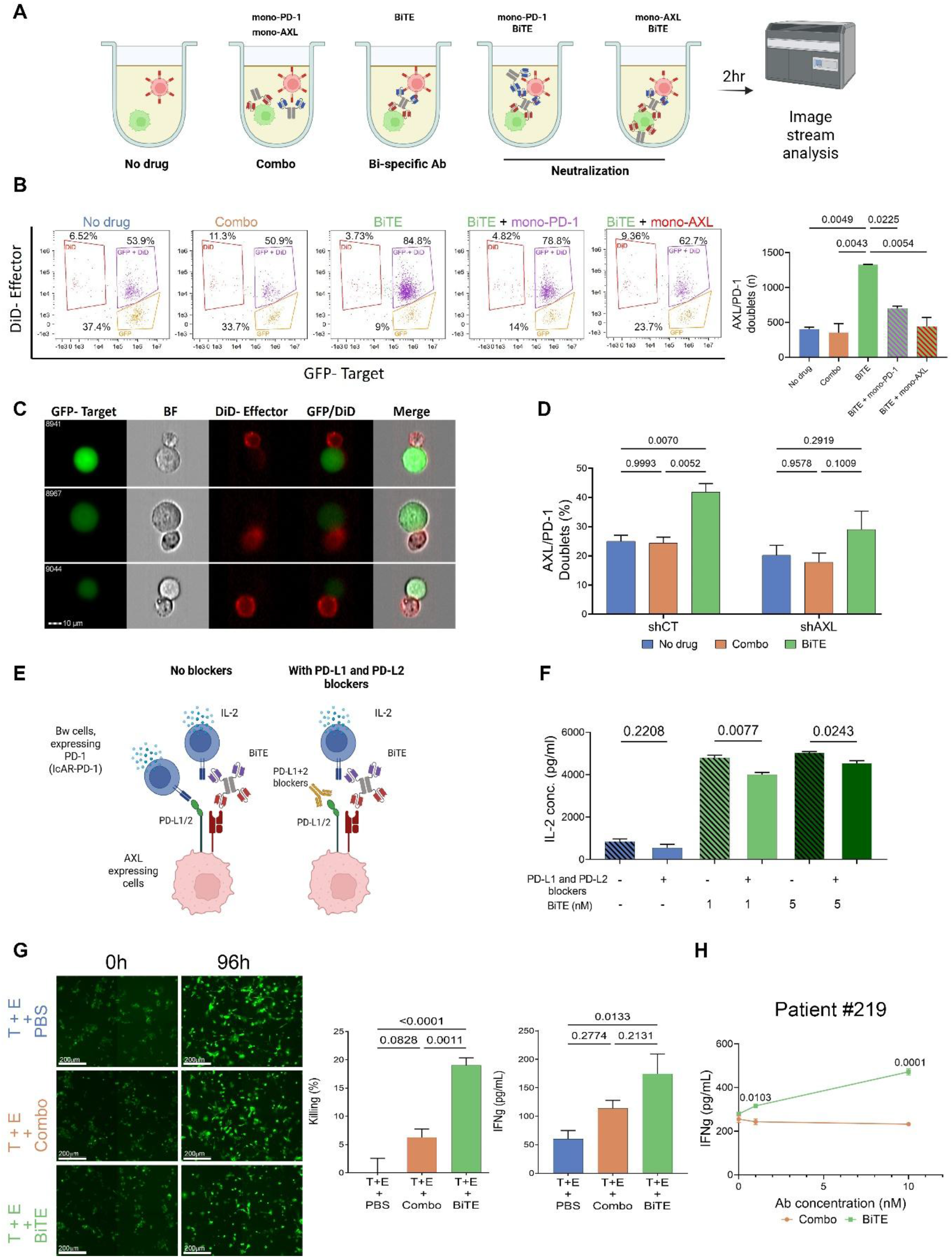
BiTE promotes tumor-T-cell interaction and enhances T-cell activation and cytotoxicity in an AXL-dependent manner. **(A)** Schematic of the doublet formation assay. GFP-labeled FaDu tumor cells and DiD-labeled IcAR-PD-1 cells (5 nM treatment). Doublets (GFP⁺DiD⁺) were quantified using ImageStream. **(B)** Left: representative dot plot showing the GFP⁺DiD⁺ population (purple gate). Right: doublet count per treatment graph. Mean ± SEM of biological duplicates (n=2). Statistical significance was determined using one-way ANOVA followed by two-tailed Tukey’s multiple comparisons test with correction for multiple comparisons (adjusted P values shown). **(C)** Representative ImageStream of BiTE-mediated immune synapse between tumor (GFP^+^) and T cells (DiD^+^). Scale bars, 10 µm. **(D)** Doublet formation assay comparing control (shCT) with AXL-knockdown (shAXL) FaDu cells with IcAR-PD-1 cells under different treatment conditions. Mean ± SEM of biological duplicates (n=2). Statistical significance was determined using two-way ANOVA followed by two-tailed Sidak’s multiple comparisons test within shCT or shAXL groups with correction for multiple comparisons (adjusted P values shown). **(E)** Schematic of the IcAR functional binding assay. FaDu and IcAR-PD-1 cells were co-cultured with BiTE, ± PD-L1/PD-L2 blockade, using IL-2 secretion as a readout. **(F)** IL-2 quantification from the functional assay. Representative of two biological experiments; mean ± SEM of technical triplicates (n = 3). Statistical significance was determined using two-tailed t-test. **(G)** Left: representative images of SCC47-GFP target cells (T) co-cultured with healthy donor PBMCs (E) at a target-to-effector (T:E) ratio of 1:10 with antibody (0.5 nM). Right: live GFP⁺ cells at 96h were quantified using JuLI Stage imaging and normalized to 0h. IFN-γ was measured by ELISA. Mean ± SEM of five technical replicates (n = 5). Statistical significance was determined using one-way ANOVA followed by two-tailed Tukey’s multiple comparisons test with correction for multiple comparisons (adjusted P values shown). Scale bar, 200µm. **(H)** IFN-γ levels from patient-derived melanoma cells #219 co-cultured with matched TILs in the presence of combination or BiTE treatment at various concentrations (0, 1 nM and 10 nM). Mean ± SEM of technical triplicates (n = 3) from single experiment. Statistical significance was determined using two-tailed t-test.

### BiTE shows favorable pharmacokinetics and tumor-targeted biodistribution, and delays tumor growth in humanized xenografts without overt toxicity

We next examined the pharmacokinetics (PK), biodistribution, safety and in vivo activity of BiTE. Bridging ELISA analyzed retention of both binding arms showed a half-life of 3.3 days after incubation of the BiTE in vitro in human serum at 37°C, as calculated by a two-compartment model (**Fig. 3A**) complete clearance occurred within 30 days (**Fig. S3A**). In vivo analysis in naïve NSG mice showed average serum half-life of 3.4 days after intraperitoneal (i.p.) injection, supporting two to three doses per week (**Fig. 3B** and **S3B**). For biodistribution, Alexa Fluor 680-conjugated BiTE was injected i.p. into NSG mice bearing AXL-expressing FaDu cell-derived xenografts (CDXs), and detected in the tumor by fluorescence imaging within 24 hours post-administration, with gradual decline in signal intensity over four days (**Fig. 3C**). At 96 hours the tumors and major organs were harvested and analyzed for fluorescent intensity (**Fig. 3D**). Tumors from BiTE-treated mice displayed significantly higher fluorescence than organs from the same mice, as well as tumors from vehicle-treated mice (**Fig. 3D**). To check for toxicity, PBMC-engrafted NSG mice dosed i.p. twice weekly with BiTE showed no significant differences in body or organ weight (**Fig. S3C** and **D**), or in markers of liver or kidney function: urea, creatinine, alkaline phosphatase (ALK-Phos), and alanine aminotransferase (ALT) (**Fig. 3E**). To examine in vivo efficacy, NSG mice bearing AXL-expressing tumors were engrafted intravenously (i.v.) with PBMCs from 3-6 healthy donors and treated i.p. with BiTE, Combo, or PBS vehicle (**Fig. 3F**). Treatment with BiTE significantly delayed subcutaneous (s.c.) SCC47 progression versus untreated (**Fig. 3G**). In an orthotopic NSG lip model, PBMCs were injected i.v. after FaDu tumor establishment, followed by bi-weekly treatments; the BiTE-treated group showed a tendency of suppressed growth versus vehicle and Combo groups (**Fig. S3E**), supporting its superior anti-tumor activity, though below statistical significance. Immunohistochemical (IHC) analysis confirmed a tendency of CD45 infiltration in the FaDu tumors treated with BiTE with significantly higher CD8 T Cells in BiTE-treated tumors compared to vehicle (**Fig. S3F**). Thus, BiTE shows significant tumor-targeted biodistribution and tumor growth inhibition in humanized xenografts without detectable systemic toxicity.

**Figure 3.**
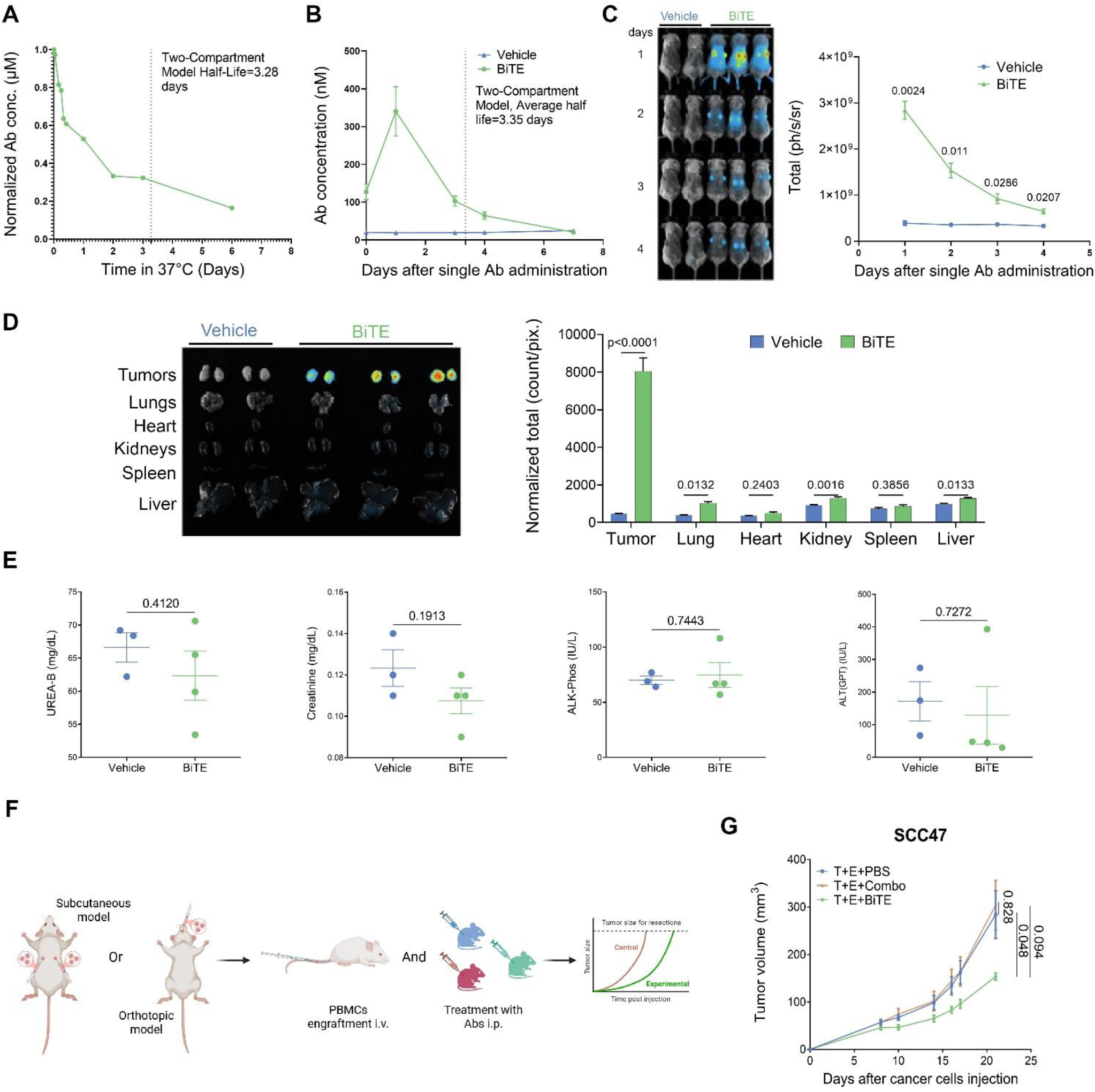
BiTE shows favorable pharmacokinetics and tumor-targeted biodistribution, and delays tumor growth in humanized xenografts without overt toxicity. (**A**) In vitro stability of BiTE during 6-day incubation at 37°C in human serum. Concentrations measured by bridging ELISA; half-life estimated using two-compartment model. (**B**) In vivo serum concentration-time profile of BiTE post-administration, quantified by bridging ELISA. Half-life estimated via two-compartment model. Mean ± SEM of four biological replicates (n=4). (**C**) In vivo time-course fluorescence imaging of NSG mice bearing FaDu cell-derived xenograft (CDX) tumors, comparing vehicle (PBS) (n=2) and BiTE (5 mg/kg) (n=3) (i.p.). Pseudo-colored signal indicates antibody distribution. Graph shows quantification across identical regions of interest (ROI). Statistical significance was determined using t-test. (**D**) Left: end-point of (C) (96 hours) fluorescence imaging of BiTE localization in tumors and organs. Right: graph shows quantification for tumors (Vehicle n=4, BiTE n=6), and organs (Kidney: Vehicle n=4, BiTE n=6, Lung/Heart/Spleen/Liver: Vehicle n=2, BiTE n=3). Statistical significance was determined using two-tailed t-test. **(E)** Serum toxicity markers assessing renal and hepatic functions. (Vehicle n=3, BiTE n=4). Statistical significance was determined using two-tailed t-test. **(F)** Schematic of the in vivo experimental design evaluating therapeutic efficacy in humanized mouse models. **(G)** Tumor volumes in subcutaneous SCC47 CDX model (T). Randomized PBMC-engrafted (E) mice received i.p. injections twice weekly: mono-Abs combination (mono-AXL+mono-PD-1, 2 mg/kg each), BiTE (2 mg/kg), or vehicle (PBS). (n = 7-8 per group). Average tumor volume ± SEM. Statistical significance was determined using a permutation test.

### Murine BiTE induces tumor growth delay in immunocompetent head and neck cancer models

To enable mechanistic studies in immunocompetent mouse models, we developed a murine BiTE and tested it in syngeneic head and neck cancer models. The murine PD-1 scFvs were designed by optimizing reported sequences (**Table S1**), while the AXL-binding scFv is cross-reactive between human and mouse[61]. We produced two murine anti-PD-1 variants (Mu-mono-PD-1 E and C), and the corresponding murine BiTEs (Mu-BiTE E and C) (**Fig. 4A**) and conducted target-binding validation by ELISA using unpurified Abs variants. Both Mu-BiTE variants displayed comparable binding to recombinant AXL and PD-1, whereas the Mu-mono-PD-1 variants showed no AXL binding, as expected (**Fig. S4A** and **B**). Based on binding, we selected variant E for large-scale production and purification of Mu-mono-PD-1 and Mu-BiTE formats (**Fig. S4C** and **D)**, hereafter referred to as Mu-mono-PD-1 and Mu-BiTE. Subsequently, we performed binding analysis of purified Abs using flow cytometry. Using murine head and neck cancer cell lines generated in our laboratory by 4-nitroquinoline-1-oxide (4NQO)-induced mutagenesis[52,53], we observed dose-dependent binding of both mono-AXL and Mu-BiTE to AXL expressed on 4NQO-T cells, with mono-AXL showing superior binding efficiency, consistent with our findings for the human Abs (**Fig. S4E**). To confirm target specificity, we generated AXL knockout in 4NQO-T cells and compared AXL binding between parental, sgCT and sgAXL cells (**Fig. S4F**). As expected, neither mono-AXL nor Mu-BiTE bound to AXL-knockout cells (4NQO-T sgAXL) (**Fig. 4B**). To assess binding to the murine PD-1 receptor, we isolated splenocytes from wild-type (WT) mice and compared Abs binding to Concanavalin A (ConA)-stimulated versus unstimulated cells. Both Mu-mono-PD-1 and Mu-BiTE bound ConA-stimulated WT splenocytes more than unstimulated cells (**Fig. 4C**). For cytotoxicity, gp100-reactive CD8⁺ T cells were co-cultured with gp100-pulsed mEERL and 4NQO-T cells, and caspase-3/7 was measured at 16 h (**Fig. 4D**). Mu-BiTE induced significantly more tumor cell death than the mono-AXL plus Mu-mono-PD-1 combination in both cell lines (**Fig. 4E** and **F**). Dose-dependent response to the Mu-BiTE was observed in an s.c. 4NQO-T CDX model in immunocompetent WT mice, with significant inhibition at 5 mg/kg (**Fig. 4G**). In an orthotopic 4NQO-T model, Mu-BiTE significantly delayed tumor progression compared to vehicle (**Fig. 4H)** and in an orthotopic lip model using the HPV-positive mEERL line, Mu-BiTE delayed tumor progression compared to both vehicle and Combo groups (**Fig. 4I**). These results show that, Mu-BiTE drives potent T-cell-mediated killing in vitro and tumor growth delay in two immunocompetent head and neck models.

**Figure 4.**
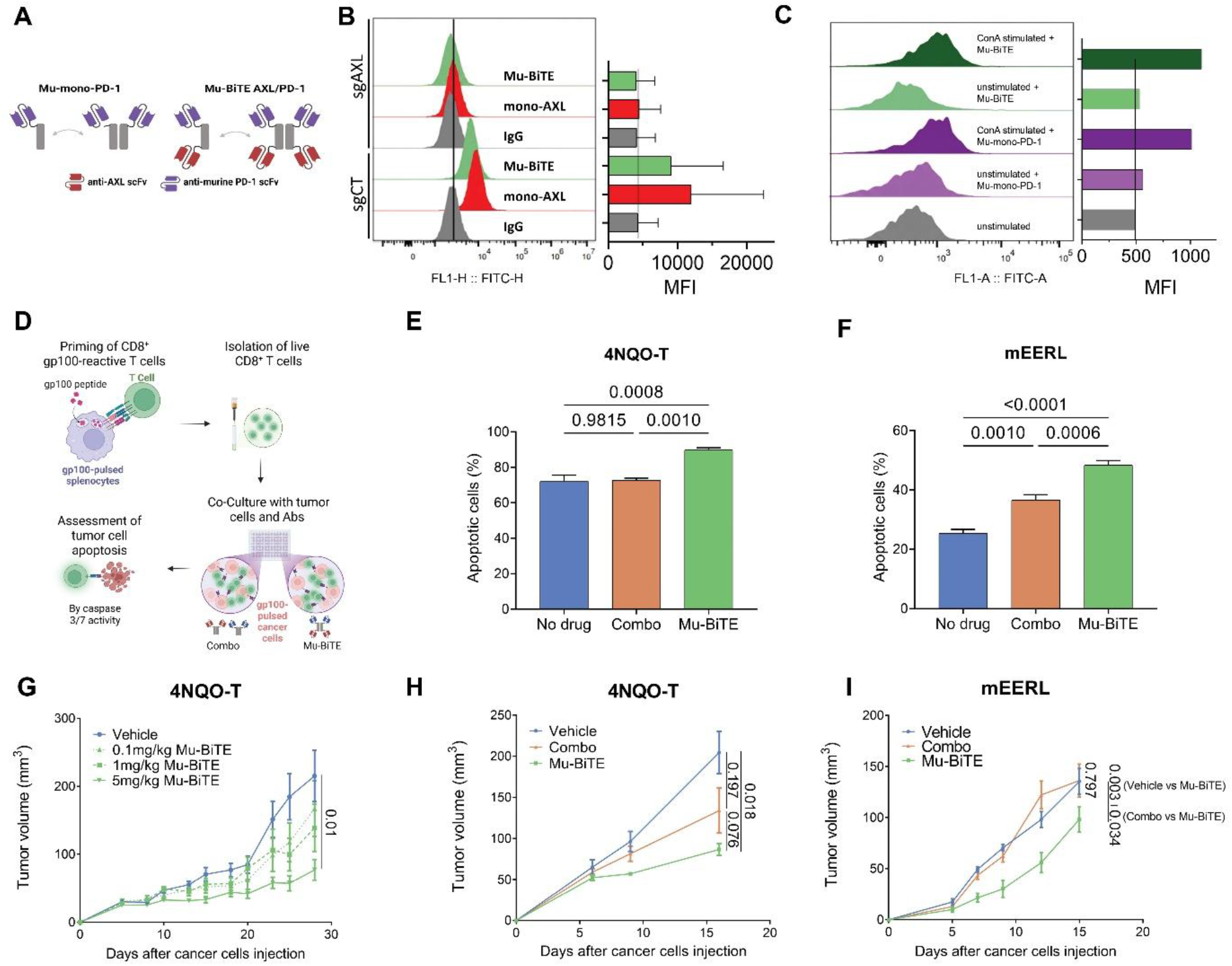
Murine BiTE induces tumor growth delay in immunocompetent head and neck cancer models. **(A)** Murine constructs binding PD-1 only (Mu-mono-PD-1) or AXL and PD-1 (Mu-BiTE-AXL/PD-1). **(B)** Binding of mono-AXL and Mu-BiTEto 4NQO-T sgCT (control) and sgAXL cells by flow cytometry. Log scale (FL1), MFI at right; representative of two biological experiments; mean ± SEM of biological duplicates (n=2). Statistical significance was determined using one-way ANOVA followed by two-tailed Tukey’s multiple comparisons test with correction for multiple comparisons (adjusted P values shown). **(C)** Binding of Mu-mono-PD-1 and Mu-BiTE to wild-type (WT) splenocytes, unstimulated versus Concanavalin A (ConA, 1 µg/mL)-stimulated by flow cytometry. Log scale (FL1), MFI at right; single experiment. **(D)** Schematic of antigen-specific cytotoxicity assay. **(E,F)** Killing of gp100-pulsed 4NQO-T (n=5) (E) and mEERL (n=4) (F) (target, T), by gp100-reactive CD8+ T cells; antibodies at 5 µg/mL, 1:4 T:E ratio; caspase-3/7 by flow cytometry at 16h. Mean ± SEM. Statistical significance was determined using a one-way ANOVA followed by two-tailed Tukey’s multiple comparisons test with correction for multiple comparisons (adjusted P values shown). **(G)** Tumor volume in 4NQO-T CDX subcutaneous model. Randomized mice received i.p. injections twice weekly: Mu-BiTE at various concentrations (0.1, 1, 5 mg/kg) or vehicle (PBS). (n = 6-10 per group). Average tumor volume ± SEM. Statistical significance was determined using a permutation test compared to vehicle only (only significant P values shown). **(H, I)** Tumor volume in 4NQO-T CDX (H) and mEERL CDX (I) orthotopic lip models. Randomized mice received i.p. injections twice weekly: mono-Abs combination (mono-AXL+Mu-mono-PD-1, 5 mg/kg each), Mu-BiTE (7.6 mg/kg molar equivalent or 5 mg/kg), or vehicle (PBS). (n = 5-7 per group). Average tumor volume ± SEM. Statistical significance was determined using a permutation test (adjusted P values shown).

### Mu-BiTE treatment reshapes the global cellular landscape across tumor zones and enhances T-cell infiltration

To assess Mu-BiTE effects on TME composition and spatial organization, we performed 18-marker (**Table S2**) multiplexed immunofluorescence on formalin-fixed, paraffin-embedded (FFPE) sections from orthotopic 4NQO-T tumors treated with vehicle, Combo or Mu-BiTE (*n=3*) (**Fig. 5A** and **B**). Cell-type composition analysis identified 11 immune/stroma cell types and two malignant cells (resting and proliferating), and comparison of cell types between the three groups (*n=3*) revealed that Mu-BiTE reorganized the TME composition relative to vehicle and Combo (**Fig. 5C**). Specifically, Mu-BiTE significantly reduced malignant epithelial cells versus vehicle (**Fig. 5D**) and proliferating epithelial cells versus both treatments (**Fig. 5E**), while DC (CD45^+^CD11c^+^) (**Fig. 5F**) and B-lymphocyte (CD45^+^B220^+^) (**Fig. 5G**) infiltration increased, significantly only for B lymphocytes (DC P = 0.09). NK (CD45^+^T-bet^+^CD161c^+^) fractions were highest in vehicle and lower in both Mu-BiTE and Combo (**Fig. 5H**), neutrophils (CD45^+^Ly6G^+^) were higher in Mu-BiTE than Combo (**Fig. S5A**), while Combo had more myeloid cells (CD45^+^CD11b^+^) (**Fig. S5B**), and endotheliocytes (CD31+) and CAFs (αSMA⁺) remained stable across treatments (**Fig. S5C** and **D**). Among T cells, Mu-BiTE increased CD4⁺ T (CD45⁺CD3⁺CD4⁺) cells versus both vehicle and Combo (**Fig. 5I**) and CD8⁺ T cells (CD45⁺CD3⁺CD8⁺) versus vehicle (**Fig. 5J**), with unsorted T cells (which can be γδ/Double-negative T cells or NKT cells), and Tregs (CD45⁺CD3⁺CD4⁺FOXP3^+^) unchanged (**Fig. S5E** and **F**). Although Mu-BiTE did not reduce exhausted CD8⁺ T cells (TOX⁺/LAG-3⁺/TIM-3⁺) (**Fig. S5G**), it markedly elevated activated (Granzyme B+) CD8⁺ T cells versus both vehicle and Combo (**Fig. 5K**). CD8⁺ T cells dominated the T-cell compartment in Mu-BiTE-treated tumors (**Fig. 5L**), whereas vehicle and Combo maintained higher CD4⁺ T cells and unsorted T lymphocytes, highlighting Mu-BiTE’s unique ability to expand and activate the CD8⁺ effector compartment.

**Figure 5.**
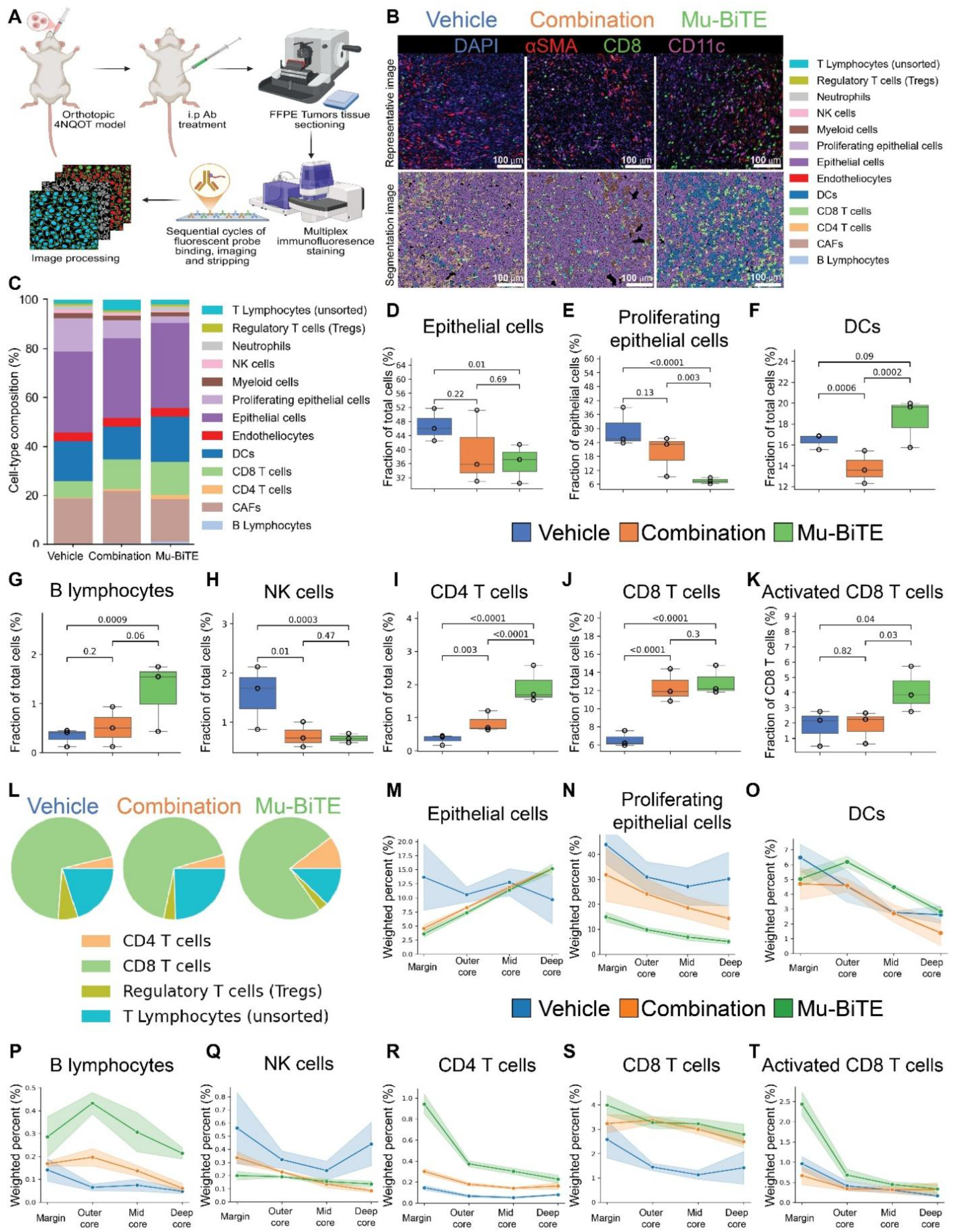
Mu-BiTE treatment reshapes the global cellular landscape across tumor zones and enhances T-cell infiltration. **(A**) Conceptual workflow for multiplex spatial data analysis. **(B)** Representative multiplex images showing nuclei (DAPI, blue), cancer-associated fibroblasts (CAFs; αSMA⁺, red), CD8⁺ T cells (CD8, green), and dendritic cells (CD11c, magenta), together with cell type–colored segmentation masks illustrating treatment-dependent remodeling of the tumor microenvironment. Scale bars, 100 µm. **(C)** Cell-type composition (%) of major cell populations across treatment groups. **(D-K)** Fractional abundance of selected cell populations across treatment groups. (D) Epithelial (tumor) cells; (E) Proliferating epithelial cells (fraction of epithelial cells); (F) DCs; (G) B lymphocytes; (H) NK cells; (I) CD4⁺ T cells; (J) CD8⁺ T cells; (K) activated CD8⁺ T cells (fraction of CD8⁺ T cells). Median and IQR with Tukey whiskers; points represent individual biological replicates (n=3). Statistical significance was determined using binomial generalized linear models with a logit link. **(L)** Composition of the T-cell compartment across treatment groups, showing the relative proportions of CD4⁺ T cells, CD8⁺ T cells, Tregs, and T lymphocytes (unsorted). **(M-T)** Zone-specific distribution of selected cell populations across treatment groups, weighted by the overall abundance of each cell type within the corresponding group. (M) Epithelial (tumor) cells; (N) Proliferating epithelial cells; (O) DCs; (P) B lymphocytes; (Q) NK cells; (R) CD4⁺ T cells; (S) CD8⁺ T cells; (T) Activated CD8⁺ T cells. Lines represent the mean and shaded bands represent the SEM of biological triplicates (n=3).

To characterize spatial organization, we segmented tumors into four concentric zones: margin, outer core, mid core, and deep core (**Fig. S5H**) and analyzed zone-specific frequencies weighted by overall cell-type abundance. The zone analysis demonstrated the unique capacity of Mu-BiTE therapy to penetrate immune-excluded areas (**Fig. S5H**). Although total malignant epithelial cells in inner zones were minimally affected by any treatment (**Fig. 5M**), Mu-BiTE maintained lower proliferating epithelial density across all zones, including the deep core (**Fig. 5N**). Unlike vehicle and Combo-treated tumors, where DC density dropped sharply from margin to core, Mu-BiTE showed a unique outer core peak and higher inner zones density (**Fig. 5O**). B cells followed similar trends (**Fig. 5P**). NK cells (**Fig. 5Q**), neutrophils (**Fig. S5I)** and myeloid cells (**Fig. S5J**) were margin-restricted, with more neutrophils overall in Mu-BiTE tumors, and more myeloid cells overall after Combo treatment, except in the deep core. Endotheliocytes remained margin-restricted (**Fig. S5K**), while CAFs were stable across zones in Combo and Mu-BiTE but margin-restricted in vehicle (**Fig. S5L**). Mu-BiTE maintained the highest CD4⁺ and CD8⁺ T-cell infiltration across all zones, with mid and deep core penetration absent in other groups (**Fig. 5R** and **S**). Unsorted T cells were evenly distributed in vehicle and Mu-BiTE but increased toward the core in Combo (**Fig. S5M**), and Tregs remained margin-localized in all groups (**Fig. S5N**), indicating that Mu-BiTE acts through effector activation rather than Treg redistribution. Critically, activated (Granzyme B^+^) CD8⁺ T cells showed margin peaks and sustained outer and mid core elevation under Mu-BiTE but remained near baseline in vehicle and Combo (**Fig. 5T**), while exhausted CD8⁺ T cells remained stable across zones and treatments (**Fig. S5O**). This sustained CD8^+^ T cells infiltration indicates that treatment with Mu-BiTE increases the activated-to-exhausted ratio rather than reducing exhaustion. In summary, while the vehicle tumors were dominated by malignant cells in the core and Combo-treated tumors by myeloid infiltration, Mu-BiTE treatment uniquely converted the tumor interiors into active zones with peak densities of DCs, B cells, and activated CD8⁺ T cells in intermediate regions.

### Mu-BiTE promotes functional immune-active spatial communities via DC-T cell engagement

To uncover relationship among malignant, stromal and immune cells, we applied spatial cellular-community analysis to determine whether these compositional changes translated into functionally relevant spatial organization. Unsupervised clustering identified 11 cellular communities (**Fig. 6A**), categorized by dominant cell-type (**Fig. 6B**). Mu-BiTE treatment significantly reduced the abundance of communities enriched in proliferating tumor cells (denoted by “pTumor” in the community names) (**Fig. 6C**): immune-desert states enriched in epithelial and proliferating epithelial cells. Mu-BiTE treatment also led to a reduction of immunosuppressive CAF-associated niches such as the pTumor-CAF-DC and Tumor-CAF (**Fig. 6D** and **E**). In contrast, Mu-BiTE tumors are enriched with Tumor-DC-CD8, CD8-DC and Tumor-DC communities composed of antigen presenting cells and activated T cells (**Fig. 6F-H**). Other communities, including Tumor, DC, CAF-DC-CD8, CAF and DC-CAF, were unchanged in total abundance (**Fig. S6A-E**), suggesting Mu-BiTE also acts through spatial redistribution and functional activation. Ranking communities by their representation in Mu-BiTE-treated tumors revealed a distinct “therapeutic signature” separating from Combo and vehicle (**Fig. 6I** and **S6F**), shifting the TME from “CAF-dominated” or “tumor-only” states to a “DC-CD8-rich” interaction state, supporting sustained anti-tumor immunity. High-resolution T-cell functional heatmaps provide mechanistic insights into how Mu-BiTE influences T-cell behavior within these communities. CD8⁺ T cells in Mu-BiTE-treated tumors concentrated in DC-related communities (**Fig. S6G**), confirming that Mu-BiTE specifically drives effector accumulation into dendritic cell-rich niches which may contribute to CD8^+^ T cells proliferation/activation. Mu-BiTE treatment increased T-cell activation across nearly every community, far exceeding Combo treatment (**Fig. 6J**). Although exhausted cells constituted a high proportion of CD8⁺ T cells in all groups given the aggressive tumor model, Mu-BiTE showed lower exhaustion in key niches such as CD8-DC and CAF-DC-CD8 (**Fig. 6K**), and a significantly higher activated-to-exhausted ratio specifically within CD8-DC and CAF-DC-CD8 niches (**Fig. 6L**). Thus, Mu-BiTE treatment efficacy reflects both increased T cell numbers and repositioning of active effectors into DC-rich hubs, whereas Combo often leaves T cells highly exhausted and poorly activated.

**Figure 6.**
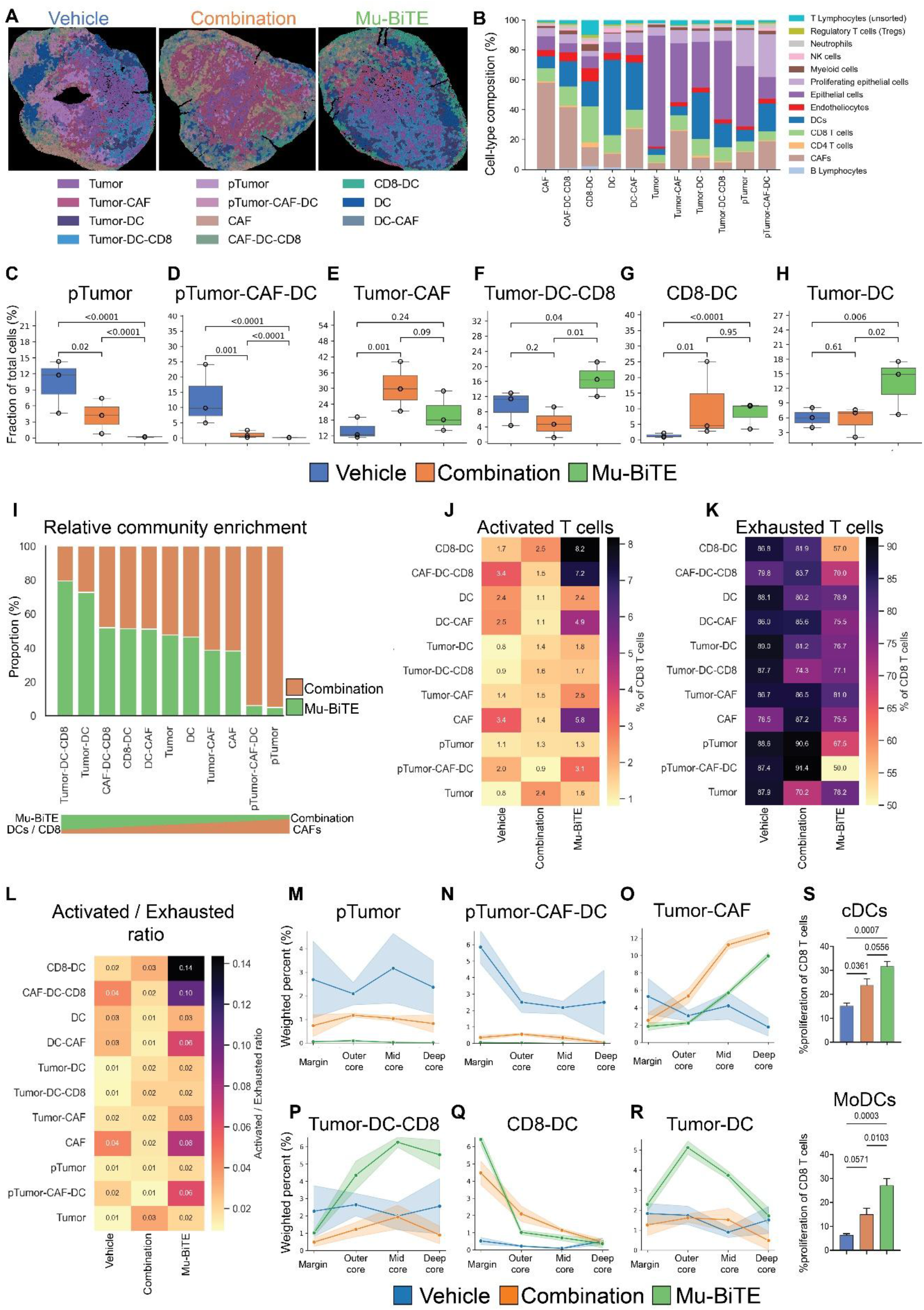
Mu-BiTE promotes functional immune-active spatial communities via DC-T cell engagement. **(A)** Representative spatial maps of cellular communities in vehicle, combination, and BiTE-treated tumors. Each cell is colored according to its community assignment, highlighting treatment-dependent changes in spatial organization, clustering, and infiltration patterns. **(B)** Normalized cellular composition of identified communities. Bars represent the mean fractional contribution of major cell types within each community. **(C-H)** Fractional abundance of selected communities across treatment groups. (C) proliferating tumor (pTumor); (D) pTumor-CAF-DC; (E) Tumor-CAF; (F) Tumor-DC-CD8; (G) CD8-DC; (H) Tumor-DC. Median and IQR with Tukey whiskers; points represent individual biological replicates (n=3). Statistical significance was determined using binomial generalized linear models with a logit link. **(I)** Relative community enrichment of Mu-BiTE and Combination tumors across communities. Bars indicate the proportion of cells contributed by each treatment group within a given community; values sum to 100% for each community. **(J,K)** Heatmaps showing the proportion of CD8⁺ T-cell functional states across communities and treatments. (J) Activated CD8⁺ T cells (Granzyme B⁺) and (K) exhausted CD8⁺ T cells, both as a percentage of total CD8⁺ T cells within each community. **(L)** Heatmap showing the ratio of activated to exhausted CD8⁺ T cells across communities and treatments. **(M-R)** Zone-specific distribution of selected communities across treatments, weighted by overall abundance within the corresponding treatment group. (M) pTumor; (N) pTumor-CAF-DC; (O) Tumor-CAF; (P) Tumor-DC-CD8; (Q) CD8-DC; (R) Tumor-DC. Lines represent the mean and shaded bands represent the SEM of biological triplicates (n=3). **(S)** Flow cytometry quantification of CD8⁺ T cell proliferation following co-culture with gp100-pulsed conventional DCs (cDCs, top) or monocyte-derived DCs (MoDCs, bottom) ± PBS / combination / BiTE treatments. Mean ± SEM of biological triplicates (n=3). Statistical significance was determined using a one-way ANOVA followed by two-tailed Tukey’s multiple comparisons test with correction for multiple comparisons (adjusted P values shown).

We then analyzed communities’ distribution across the four tumor zones (weighted by overall abundance). Mu-BiTE treatment reorganized the tumor interior, moving immune activity from periphery to core and restricting or reversing suppressive, immune-desert communities (**Fig. S6H**). Vehicle showed high pTumor distribution peaking in mid-core, whereas Mu-BiTE treatment reduced this desert state across all zones to the lowest of any group (**Fig. 6M**). The suppressive pTumor-CAF-DC niche was prominent and mid-core-peaking in vehicle but was decreased in inner zones and restricted to the margin under Mu-BiTE (**Fig. 6N**). While both Combo and Mu-BiTE tumors showed an increasing Tumor-CAF distribution trend, with density rising from the outer core and peaking in the deep core, Mu-BiTE maintained lower distribution across all zones, particularly in outer and mid core where immune-active communities were simultaneously enhanced (**Fig. 6O**). In contrast, immune-active communities expanded into previously excluded regions: the Tumor-DC-CD8 community peaked in mid-core specifically in Mu-BiTE-treated tumors while remaining low and flat in vehicle and Combo tumors (**Fig. 6P**); indicating complex immune interactions were successfully organized deep within the tumor mass. The CD8-DC community was margin-dominant in Combo and Mu-BiTE tumors and very low in vehicle tumors (**Fig. 6Q**). The Tumor-DC community showed the most striking shift, rising from margin to a mid- and deep-core peak in Mu-BiTE tumors while flat and low in other groups (**Fig. 6R**), suggesting that Mu-BiTE uniquely facilitates dendritic cell engagement with malignant cells in territories where these interactions are normally absent. The tumor community trended upward toward the deep core in Mu-BiTE and Combo tumors but was lower and more stable in vehicle (**Fig. S6I**). The remaining communities, DC, CAF-DC-CD8, CAF, and DC-CAF, were largely treatment-independent, with mostly margin-dominant distributions reflecting tumor architecture, except CAFs which were evenly distributed in Combo and Mu-BiTE tumors (**Fig. S6J-M**). To test our in vivo observation that and Mu-BiTE increases CD8^+^/DC interaction, we co-cultured CD8⁺ T cells with gp100-pulsed conventional DCs (cDCs) or monocyte-derived DCs (MoDCs) and treated with Mu-BiTE, Combo or PBS. Flow cytometry analysis showed that when interacting with gp100-pulsed DCs, Mu-BiTE significantly increased CD8⁺ T-cell proliferation compared with Combo and control (**Fig. 6S**), supporting the in vivo spatial findings and indicating that Mu-BiTE-mediated DC interactions contribute to T-cell expansion within the tumor. Collectively, these results attribute the superior efficacy of Mu-BiTE therapy to conversion of immune-excluded tumor territories into active hubs of DC-T cell engagement.

## Discussion

PD-1 immune checkpoint blockade is a central cancer therapy, yet expression of AXL limits its efficacy by impairing tumor cell killing and fostering an immunosuppressive TME[16,37,62]. Preclinical studies show that AXL inhibition enhances PD-1 blockade, an approach currently under clinical evaluation[37,63–66]. However, conventional combination therapy with separate anti-AXL and anti-PD-1 blockers lacks spatial coordination. These agents circulate independently and do not ensure physical proximity between PD-1^+^ effector T cells and AXL-expressing immunosuppressive cells within the heterogeneous TME, potentially limiting synergy where AXL-driven immune suppression is most active. We therefore developed anti-AXL, anti-PD-1 Fc-scFvs, alongside a tetravalent scFv BiTE and compared combination treatment (Combo) with the BiTE. Tetravalent formats provide increased avidity and sustained target engagement relative to bivalent formats, therefore reducing antigen escape, and have been associated with improved efficacy and safety[67–69]. These properties may improve immune engagement, tumor targeting, and therapeutic outcomes, suggesting advantages over bivalent formats[45]. We incorporated a mutated Fc to extend half-life over scFv-only BiTEs. Screening three BiTE configurations, two with tandem and one non-tandem scFvs, by PD-1-dependent IL-2 secretion, prioritizing function over affinity alone, identified the non-tandem BiTE as the most active and optimized for T-cell engagement. Biochemical characterization showed that the BiTE retained moderate binding to AXL and high-affinity to PD-1 with relatively slow dissociation, indicating stable receptor engagement. Importantly, the BiTE did not simply reproduce the effects of Combo treatment but generated a distinct outcome: AXL-dependent tumor-T-cell conjugation, enhanced activation, increased killing, and improved tumor control, indicating that spatial coordination between PD-1⁺ T cells and AXL-expressing cells is central to this mechanism, rather than simultaneous blockade alone. Although the BiTE bound AXL with lower affinity than mono-AXL, it retained specific binding and function, indicating that dual engagement and enforced cell-cell contact can compensate for reduced single-arm affinity, consistent with its superiority over Combo in vitro and in vivo. Whereas most bispecific T-cell engagers recruit T cells through CD3e[44], including a recent AXL-CD3 bispecific T-cell engager in ovarian cancer[70], we chose to engage PD-1 for several reasons. First, AXL-expressing tumors, such as those with partial epithelial-mesenchymal transition (pEMT), are immune-excluded, and their tumor-resident T cells frequently express PD-1[71–74]. Thus, targeting PD-1 directly engages these cells, enhances activation and may limit progression toward functional exhaustion, resulting in more effective tumor cell killing, as our data show. Second, CD3 engagement risks non-specific polyclonal activation and off-target toxicity[44], whereas the BiTE preferentially activates T cells that are already within the TME, restricting immune activation to the tumor site and reducing off-target effects. Third, BiTE-mediated killing was achieved without CD3e engagement, further limiting systemic toxicity. Fourth, AXL-expressing cells also express PD-L1 and PD-L2[17,22,75] which suppress T-cell mediated killing, so blocking the PD-1/PD-L1 axis is critical for durable anti-tumor activity. Finally, AXL is expressed across multiple TME compartments, including malignant cells[26,27], TAMs[28–30], and DCs[28,31]. Following BiTE treatment, CAF was reduced and DCs increased, changes associated with enhanced T-cell proliferation and immune activity within the tumor. However, spatial analysis suggests that the CAF effect should be interpreted primarily as reduced CAF-associated suppressive niches rather than global CAF depletion. This distinction suggests that the BiTE reorganizes stromal-immune interactions rather than simply eliminating stromal populations. Thus, dual AXL/PD-1 therapy facilitates direct tumor killing and reprograms the tumor stroma toward a more permissive immune environment. Moreover, enhanced T-cell-DC interaction mediated by BiTE supports sustained T-cell proliferation and activation, a mechanism reported for DC/CD3 bispecific T-cell engagers [76,77]. This multi-compartment effect may offer an advantage over CD3-based bispecific T-cell engagers which rely on broad T-cell activation. Since non-tandem scFv Ab configuration can significantly affect biological activity[78,79], we used a functional quantitative PD-1 assay[51] and found that the non-tandem tetravalent configuration had superior potency, likely due to enhanced stability and ligand/scFv binding. A mutated Fc domain[80] gave a serum half-life of approximately 3.5 days; together with preferential tumor accumulation and absence of weight or liver/kidney changes in PBMC-humanized NSG mice, this indicates local action and tolerability[81]. Efficacy studies in PBMC-humanized NSG mice revealed significant tumor growth inhibition, although the magnitude of response was limited by low human T-cell infiltration, a known limitation of this model[82]. We therefore used immunocompetent models, in which the murine BiTE delayed tumor growth more effectively than Combo in two orthotopic head and neck models, indicating dual AXL/PD-1 engagement contributes to optimal activity. A central finding of this study is that the Mu-BiTE reorganizes the TME spatial architecture through coordinated cellular redistribution and functional reprogramming. It increased CD8⁺ T-cell infiltration and activated CD8⁺ T cells abundance, enriched DCs, reduced proliferating epithelial cells, and favored DC-CD8 community over CAF-dominated niches relative to Combo treatment. These findings are consistent with clinical observations in HNSCC, where CAF- and myeloid-enriched tumors are associated with poor prognosis, while immune-infiltrated tumors exhibit improved outcomes[83]. While it is unclear why Combo treatment induced or maintained CAF-associated communities, these CAF-rich niches may block CD8 T-cell activity as recently shown in HNSCC patients[84]. Thus, the superiority of the BiTE likely reflects not only enhanced killing but also redirection of immune cells into productive DC- and CD8-rich niches. This spatial reprogramming explains why the BiTE was more effective than conventional monospecific Combo therapy.

Several limitations of this study should be acknowledged. First, a format-matched tetravalent control with one inactive arm was not included, so spatially enforced dual engagement cannot be fully disentangled from increased tetravalent avidity. Second, although our data demonstrates a strong association between BiTE treatment, enhanced T-cell activation, and spatial TME reorganization, the precise mechanism by which physical interaction enhances T-cell activation and proliferation was not investigated in this study. Therefore, additional functional experiments are needed to determine whether enrichment of DC-CD8 and tumor-DC-CD8 communities directly drives the improved anti-tumor response or follows tumor-lysis that promotes DC accumulation and co-localization with CD8 T cells. Third, since AXL is expressed across several TME compartments, it remains unclear whether malignant-cell AXL expression is required as a biomarker for response to BiTE. Fourth, the causal necessity of CD8 T cells, DCs, and compartment-specific AXL expression was not established and requires in vivo depletion and cell-type-specific AXL deletion. Finally, the PBMC-humanized NSG model only partially recapitulates the human immune system and shows limited human T-cell infiltration, which may have restricted the magnitude of response. Although Mu-BiTE allowed evaluation in immunocompetent models, differences between murine and human immune systems, Ab affinity, Fc behavior, and receptor biology may affect translatability.

## Conclusions

In conclusion, the anti-AXL/PD-1 BiTE provides a therapeutic advantage over conventional anti-AXL plus anti-PD-1 combination therapy by enforcing spatially coordinated engagement between PD-1⁺ T cells and AXL-expressing tumor and DC cells, enhancing T-cell activation and cytotoxicity, reshaping the TME toward immune-active DC-CD8-rich communities, and potentially overcoming the functional and spatial barriers that limit PD-1 blockade in AXL-expressing tumors.

## Abbreviations

4NQO: 4-nitroquinoline-1-oxide
Abs: Antibodies
ADCC: Ab-Dependent Cellular Cytotoxicity
ALK-Phos: Alkaline phosphatase
ALT: Alanine aminotransferase
BiTE: Bispecific T-cell engagers
BSA: Bovine serum albumin
CAF: Cancer associated fibroblasts
CDC: Complement-dependent cytotoxicity
cDCs: Conventional dendritic cells
CDX: Cell-derived xenografts
ConA: Concanavalin A
DCs: Dendritic cells
ELISA: Enzyme-linked immunosorbent assay
FBS: Fetal bovine serum
FFPE: Formalin-fixed, paraffin-embedded
HNSCC: Head and neck squamous cell carcinoma
i.p: Intraperitoneal
i.v: Intravenously
IFN: Interferon
IHC: Immunohistochemical
IL: Interleukin
MoDCs: Monocyte-derived dendritic cells
PBMCs: Peripheral blood mononuclear cells
PBS-T: Phosphate-buffered saline Tween
PD-1: Programmed cell death protein 1
PD-L1/2: Programmed death-ligand 1/2
pEMT: Partial epithelial-mesenchymal transition
PFA: Paraformaldehyde
PK: Pharmacokinetics
s.c.: Subcutaneous
scFvs: Single-chain variable fragments
SEC-HPLC: Size-exclusion high-performance liquid chromatograph
SPR: Surface plasmon resonance
TAMs: Tumor-associated macrophages
TILs: Tumor-infiltrating lymphocytes
TME: Tumor microenvironment
WT: Wild-type

## Declarations

### Ethics approval and consent to participate

Animal experiments were conducted in compliance with protocols established by the Institutional Animal Care and Use Committee (IACUC) of Ben-Gurion University of the Negev for ensuring animal welfare and minimizing discomfort. The animal ethical clearance protocol number used for the study were protocols IL-44-06-2019E and IL-64-10-2023E. Patients with metastatic melanoma who failed on anti-PD1 were recruited by the Ella Lemelbaum Institute for immuno-oncology and melanoma according to approval no. SMC-3566-16 approved by the Sheba Medical Center Ethics committee.

### Consent for publication

Not applicable.

### Data availability

The data supporting the findings of this study are available within the article and its Supplementary Information. The raw data underlying the graphs and the flow cytometry, ELISA, SPR, and western blot experiments shown in the figures are available from the corresponding author upon request. The multiplex spatial-imaging (CODEX/PhenoCycler) dataset generated in this study is available from the corresponding author upon request. The scFv sequences generated for this study are provided in Supplementary Table S1. Any additional data supporting the findings of this study are available from the corresponding author upon request. All data supporting the findings of this study, including those currently available upon request, will be made freely available without restriction upon publication of the manuscript.

### Code availability

The custom code used to process and analyze the multiplex spatial-imaging (CODEX/PhenoCycler) data in this study is available from the corresponding author upon request, and will be deposited in a publicly accessible repository (e.g., Zenodo or GitHub) upon acceptance of the manuscript.

### Competing interests

The authors declare that they have no competing interests.

## Funding

This work was funded by Israel Cancer Research Fund acceleration (ICRF, 81071411 to A.P. and M.E.); Israel Science Foundation (ISF, 302/21 to M.E.); Ministry of Innovation, Science & Technology (MOST #5912 to M.E. and O.C.); NSFC-ISF joint research program (M.E and D.K #3409/20); NSFC-ISF joint research program (M.E and H.W 2929/23).

To O.C.: work was supported by the Israel Science Foundation (ISF, 2802/22), the United States-Israel Binational Science Foundation (BSF, 2023129) and the Israel Cancer Research Fund (ICRF, 81233611).

## Authors’ Contributions

Conceptualization, A.A and M.E; Methodology, A.P; Investigation, M.C and M.E; Resources, R.S.F, D.K, H.W, A.P and L.K; Designed, K.M.Y, O.I, Y.W, A.A and M.E; Performed, K.M.Y, B.K, D.R, D.S.M, A.K, R.B.U, O.L.G, I.M, A.R, A.V, A.N, N.Z, D.B.M, E.R, O.N, J.Z, S.A.E, S.J, G.E, K.W, R.A.S, M.W, U.H, O.I, Y.W and Y.C; Spatial analysis, G.P, L.K and O.C; Wrote, K.M.Y, T.S, G.P, O.C and M.E; Supervision, O.C and M.E

## Acknowledgements

The authors gratefully acknowledge the support of the Jules Harris Chair in Oncology, held by Prof. Moshe Elkabets.

Optical imaging data was acquired at the de Picciotto Cancer Cell Observatory, in memory of Wolfgang and Ruth Lesser, of the Moross Integrated Cancer Center in the Department of Life Sciences Core Facilities at the Weizmann Institute of Science.

Figures 1-5 and Supplementary Figure S1 contain illustrations created with BioRender.com.

## Notes

### Competing Interest Statement

The authors have declared no competing interest.

