## Supplementary figures and tables for "Spatially enforced engagement of PD-1⁺ T cells with AXL⁺ cells reprograms the tumor microenvironment and overcomes anti-PD-1 resistance"

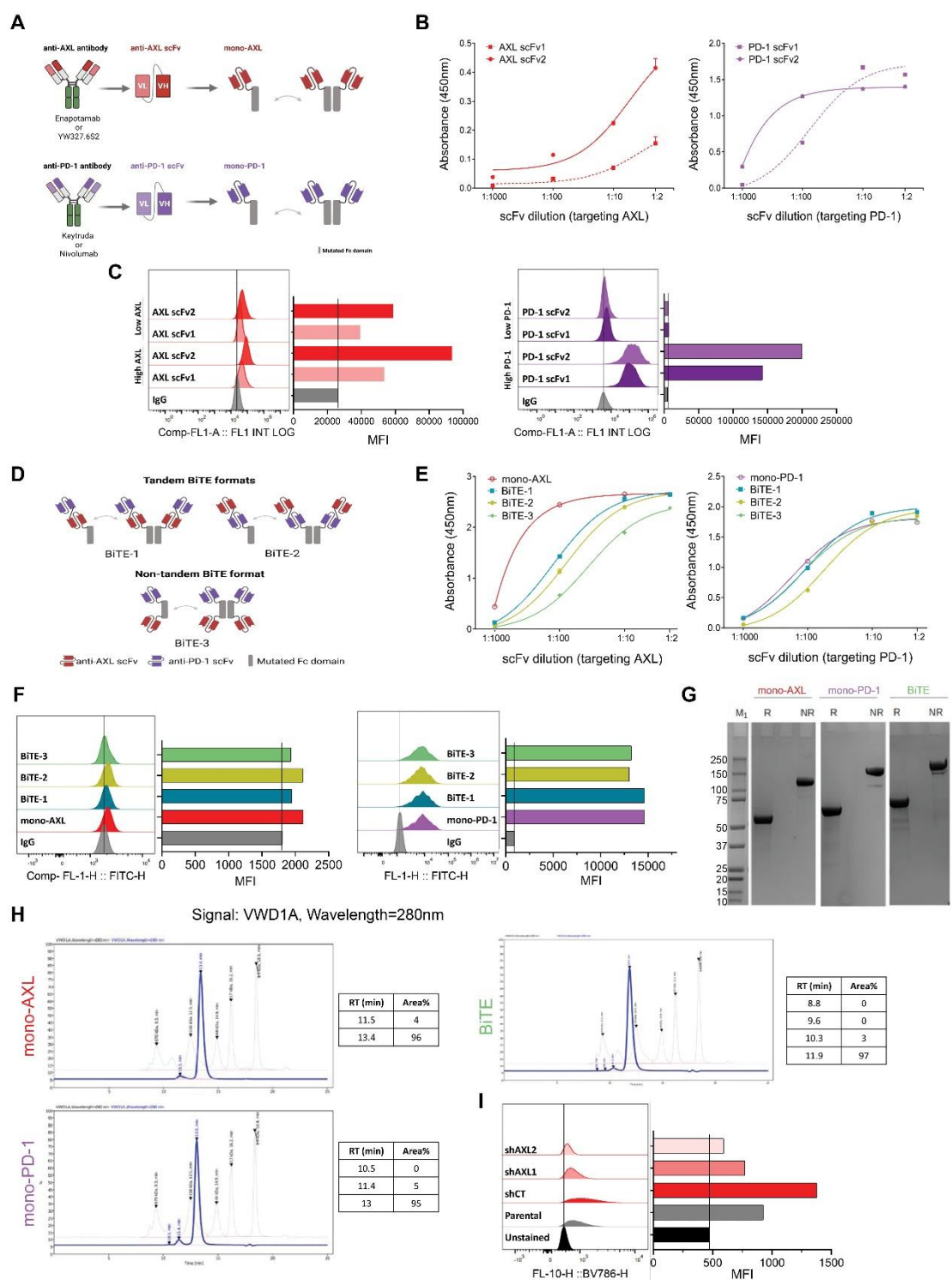

**Figure S1. Development and characterization of a non-tandem tetravalent bispecific antibody targeting AXL and PD-1.**

(A) Design of monospecific antibodies targeting AXL and PD-1, built from known scFvs fused to a mutated Fc domain. (B) ELISA detecting the binding of mono-specific scFv variants to recombinant AXL or PD-1. Mean ± SEM of technical triplicates ($n = 3$) from single experiment. (C) Flow cytometry of anti-AXL (AXL scFv1/2) and anti-PD-1 (PD-1 scFv1/2) variants on cells with differential target expression (AXL: SCC47 shCT vs shAXL; PD-1: IcAR-PD-1 vs IcAR-WT). Log scale (FL1), MFI at

right; single experiment. **(D)** Design of BiTE configurations. **(E)** ELISA binding of BiTE variants to AXL (left) and PD-1 (right). Mean ± SEM of technical triplicates ($n = 3$) from single experiment. **(F)** Flow cytometry of BiTE variants as in C. Log scale (FL1), MFI at right; single experiment. **(G)** SDS-PAGE analysis of mono-AXL, mono-PD-1, and BiTE under reduced (R) and non-reduced (NR) conditions. **(H)** Chromatographic analysis of mono-AXL, mono-PD-1, and BiTE by SEC-HPLC at 280 nm. RT, retention time. **(I)** Flow cytometry of AXL on AXL knockdown FaDu cells. Log scale (FL10), MFI at right; single experiment.

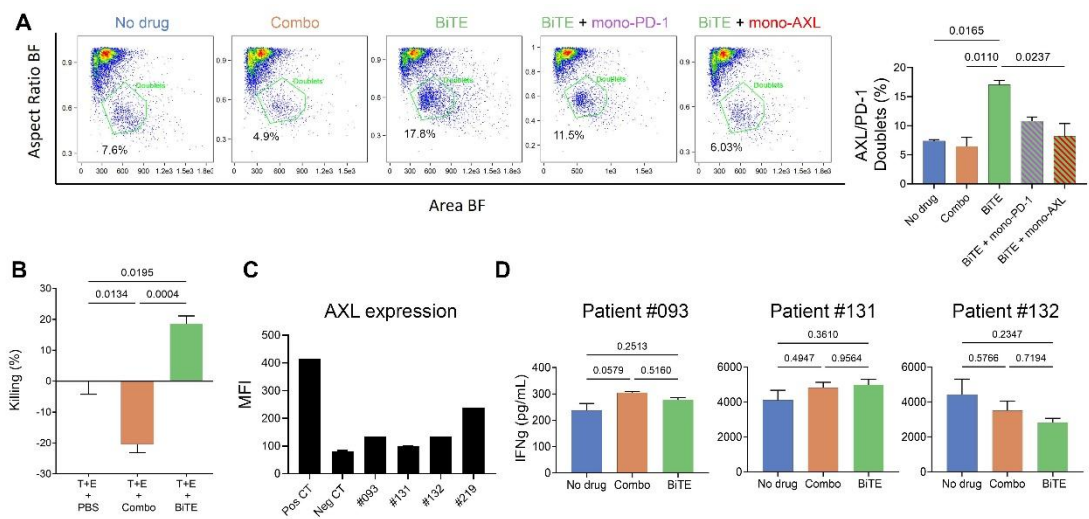

**Figure S2. BiTE promotes tumor-T-cell interaction and enhances T-cell activation and cytotoxicity in an AXL-dependent manner.** **(A)** Left: co-incubated GFP-labeled FaDu cells and DiD-labeled IcAR-PD-1 cells were treated with BiTE-AXL/PD-1, mono-antibody combination, BiTE + individual mono-antibodies, or no drug. Right: dot plot graphs of Area vs. Aspect Ratio (brightfield) identify total doublet percentages. Representative of two biological experiments; mean ± SEM of biological duplicates ($n=2$). Statistical significance was determined using a one-way ANOVA followed by two-tailed Tukey's multiple comparisons test with correction for multiple comparisons (adjusted P values shown). **(B)** Percentage of cell killing following co-culture of FaDu target (T) cells and PBMC effectors (E) under combination, BiTE treatment or with no drug. Mean ± SEM of technical replicates (T+E - $n = 2$, Combo, BiTE $n=3$) from single experiment. Statistical significance was determined using a one-way ANOVA followed by two-tailed Tukey's multiple comparisons test with correction for multiple comparisons (adjusted P values shown). **(C)** Flow cytometry analysis of AXL expression in control and patient-derived samples shown by geometric mean fluorescence intensity. Single experiment. **(D)** Supernatant IFN-γ secretion measured by ELISA from killing assays using patient-derived tumor samples #093, #131, and #132 as target cells and their corresponding matched TILs as effector cells treated with vehicle/no treatment, combination treatment, or BiTE alone. (10:1 E:T ratio; 10 nM antibody treatments). Mean ± SEM of technical triplicates ($n = 3$) from single experiment.

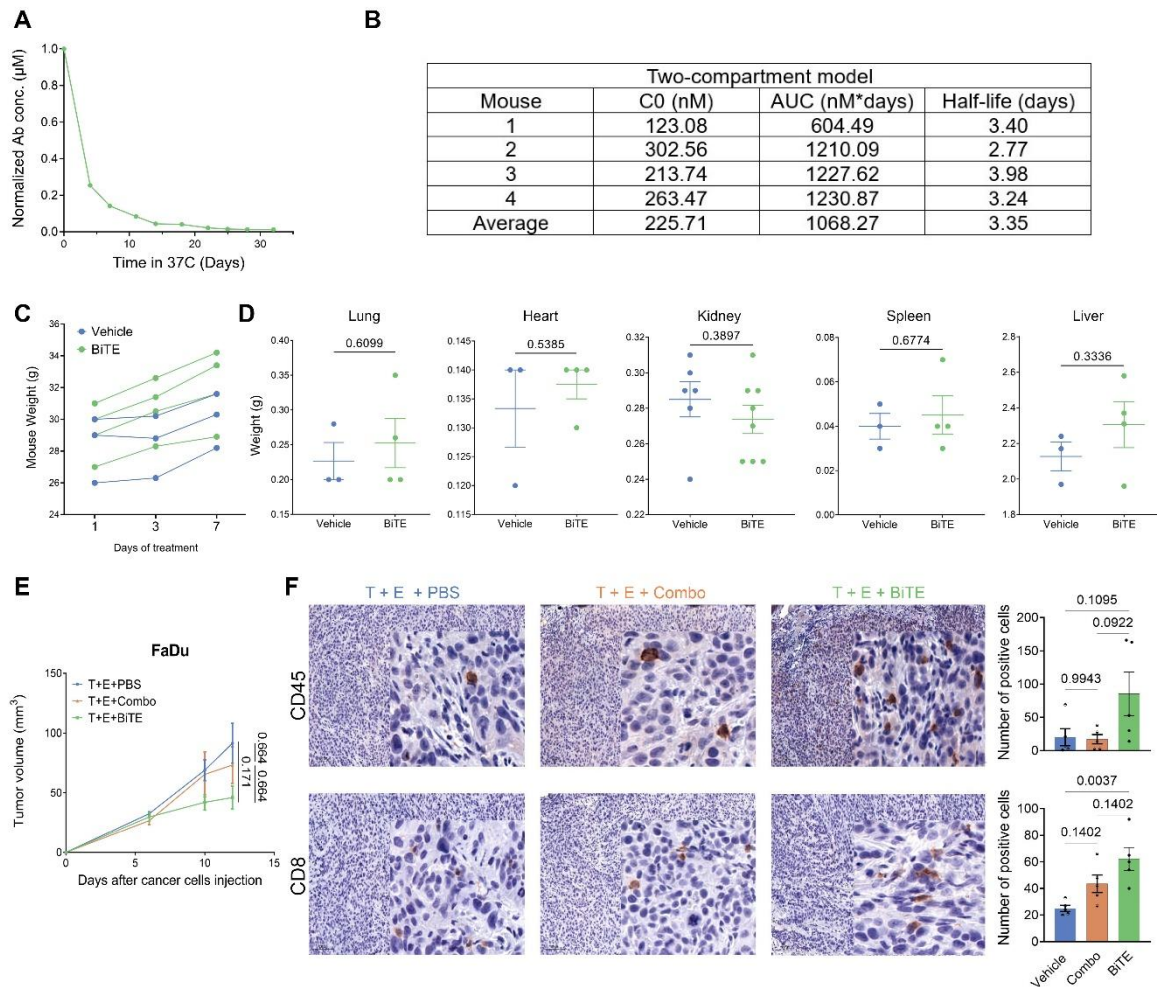

**Figure S3. Stability, pharmacokinetics, and tolerability of BiTE, with tumor growth and immune infiltration.** (A) In vitro stability of BiTE during 1-month incubation at 37°C in human serum. Concentrations measured by bridging ELISA. (B) Estimated pharmacokinetic parameters for individual mice following administration of BiTE in vivo. The table shows the initial concentration ($C_0$, nM), area under the concentration-time curve (AUC, nM·days), and elimination half-life ($t_{1/2}$, days) estimated using two-compartment model. Mean values across four mice are reported in the final row. (C) Body weight changes over time during the toxicity study in mice treated with vehicle control (n=3) or BiTE (n=4). (D) Endpoint comparison of internal organ weights (Kidney - Vehicle n=6, BiTE n=8, Lung, Heart, Spleen and Liver - Vehicle n=3, BiTE n=4) from mice treated with vehicle control or BiTE. Statistical significance was determined using two-tailed t-test. (E) Tumor volume measurements in FaDu CDX orthotopic lip model (T). Randomized PBMC-engrafted (E) mice received i.p. injections twice weekly: mono-Abs combination (mono-AXL+mono-PD-1, 5 mg/kg each), BiTE (7.6 mg/kg molar equivalent), or vehicle (PBS). (n = 5 per group). Average tumor volume ± SEM. Statistical significance was determined using a permutation test. (F) IHC representative images and quantification of CD45 and CD8 staining in FaDu Tumors at experiment end point.

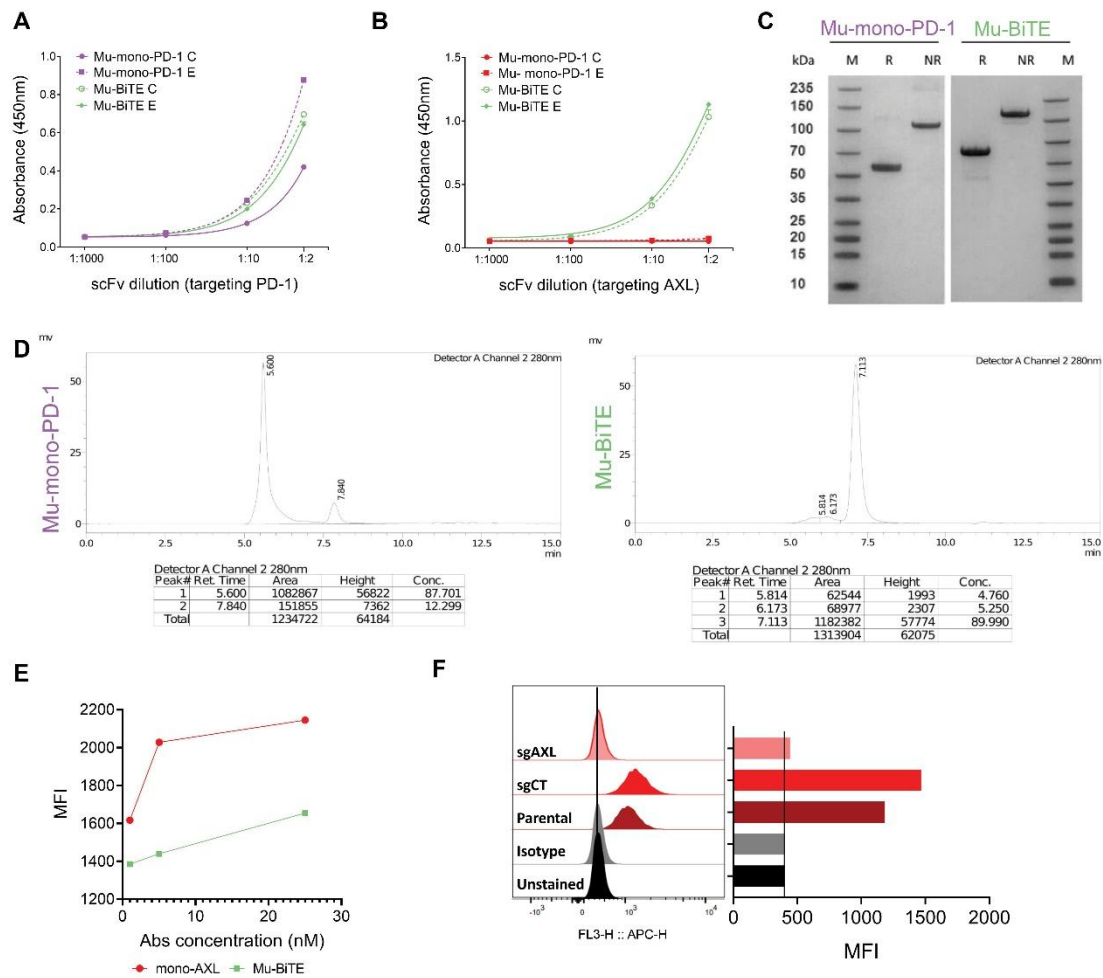

**Figure S4. Murine BiTE constructs: generation and in vitro characterization.** (A, B) ELISA detecting the binding of Mu-mono-PD-1 and Mu-BiTE variants to recombinant PD-1 (A) and AXL (B). Mean ± SEM of technical triplicates ($n = 3$) from single experiment. (C) SDS-PAGE analysis of Mu-mono-PD-1 and Mu-BiTE under R and NR conditions. (D) Chromatographic analysis of Mu-mono-PD-1 and Mu-BiTE detected by UV absorbance at 280 nm. Peak retention times and relative peak percentages are summarized below each chromatogram. (E) Binding curves of mono-AXL and Mu-BiTE to 4NQO-T cells across increasing concentrations (Ab concentration, nM) measured by flow cytometry for mean fluorescence intensity (MFI); single experiment. (F) AXL expression on 4NQO-T parental, sgCT, and sgAXL cells were detected using anti-AXL APC Ab by flow cytometry. Log scale (FL3), MFI at right; single experiment.

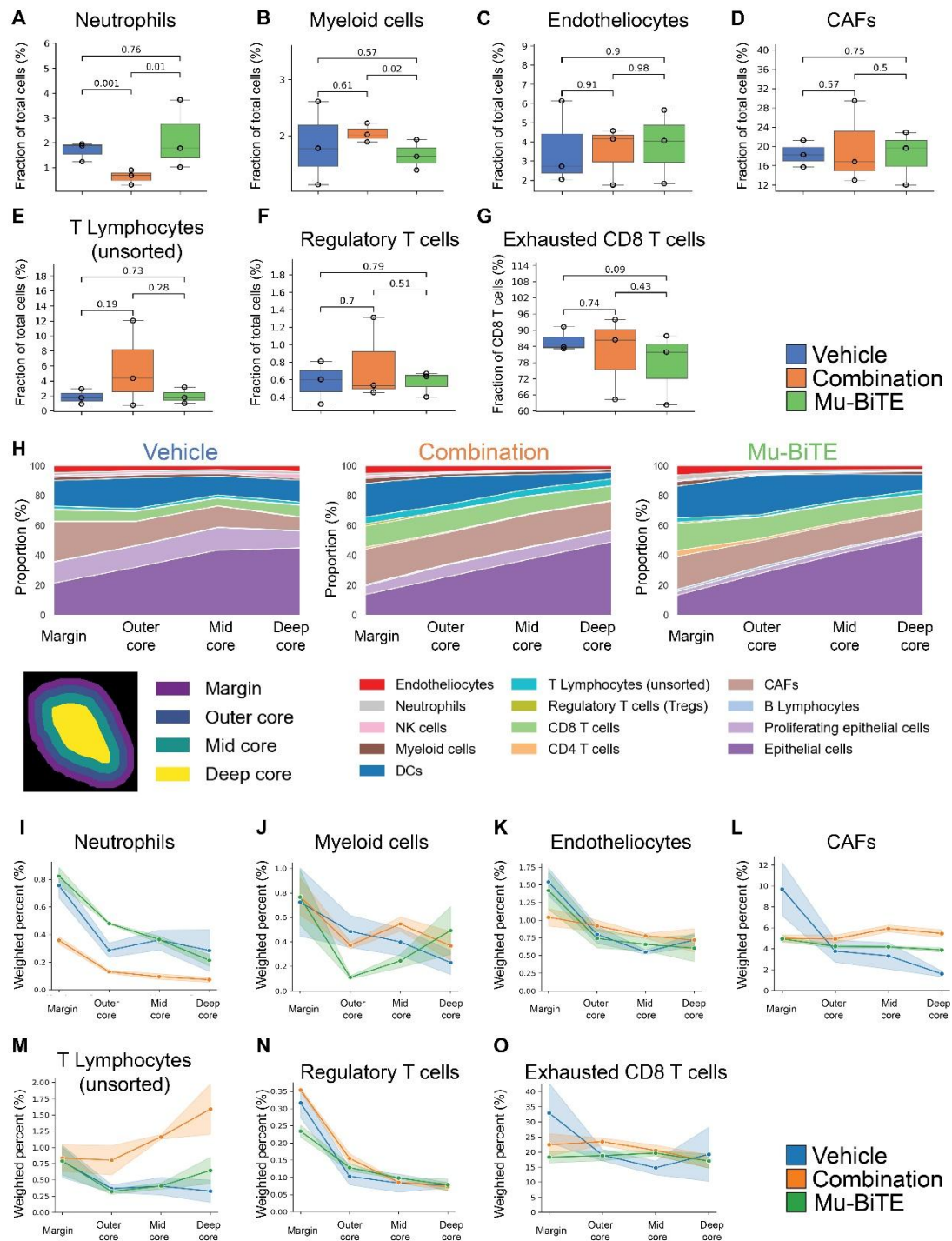

**Figure S5. Abundance and zonal distribution of additional cell populations across treatment groups.** (A-G) Fractional abundance of selected cell populations across treatment groups. (A) Neutrophils; (B) Myeloid cells; (C) Endotheliocytes; (D) Cancer-associated fibroblasts (CAFs); (E) T lymphocytes (unsorted); (F) Regulatory T cells (Tregs); (G) Exhausted CD8<sup>+</sup> T cells (fraction of CD8<sup>+</sup> T cells). Median and IQR with Tukey whiskers; points represent individual biological replicates (n=3). Statistical significance was determined using binomial generalized linear models with a logit link. (H) Relative cellular composition of tumor zones. Stacked area plots show the proportion of individual cell populations within the Margin, Outer

Core, Mid Core, and Deep Core zones, with each zone normalized to 100%. **(I-O)** Zone-specific distribution of selected cell populations across treatment groups, weighted by the overall abundance of each cell type within the corresponding treatment group. (I) Neutrophils; (J) Myeloid cells; (K) Endotheliocytes; (L) Cancer-associated fibroblasts (CAFs); (M) T lymphocytes (unsorted); (N) Regulatory T cells (Tregs); (O) Exhausted CD8<sup>+</sup> T cells. Lines represent the mean and shaded bands represent the SEM of biological triplicates (n=3).

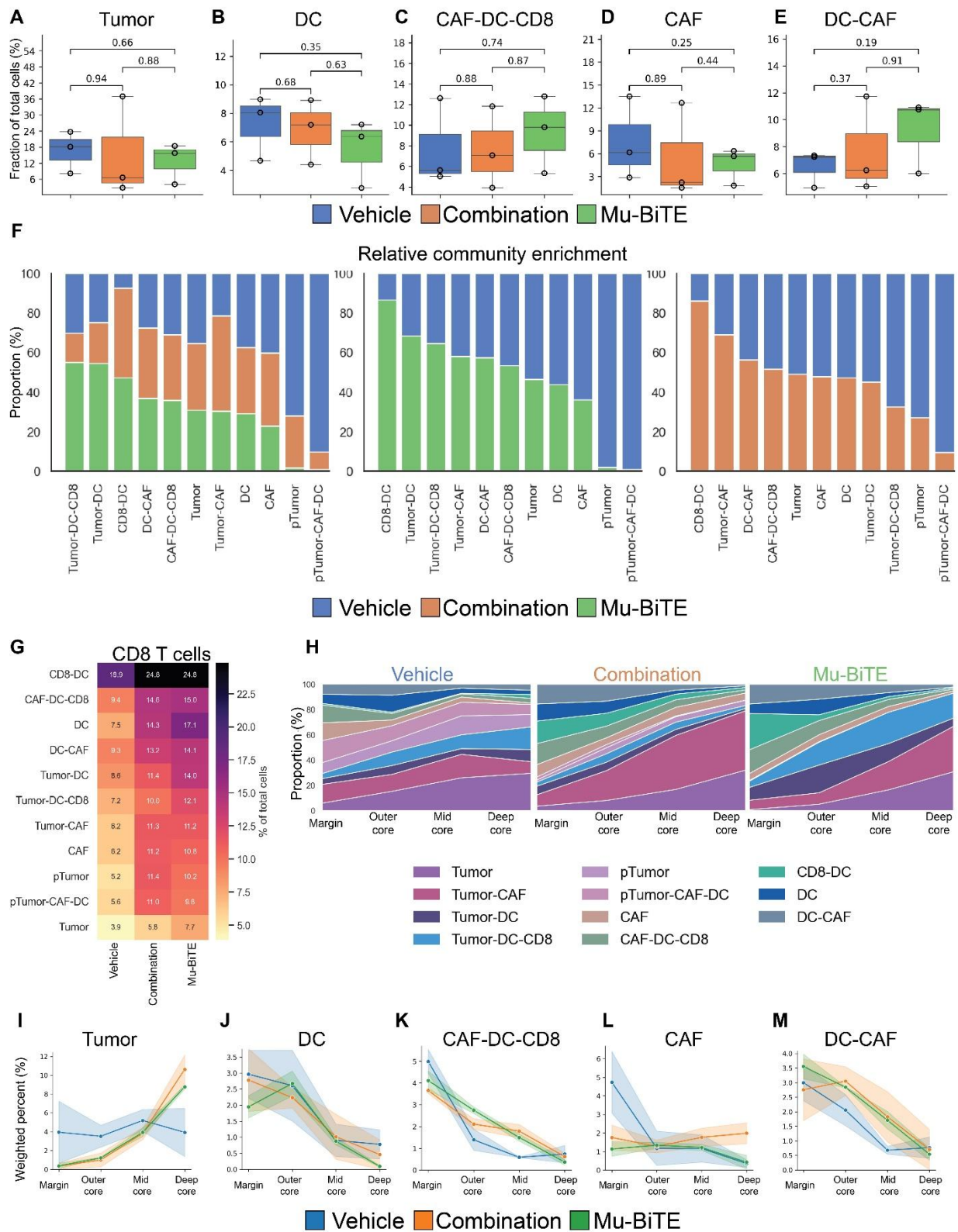

**Figure S6. Abundance, enrichment, and zonal distribution of spatial cellular communities across treatment groups. (A-E)** Fractional abundance of selected communities across treatment groups. (A) Tumor; (B) DC; (C) CAF-DC-CD8; (D) CAF; (E) DC-CAF. Median and IQR with Tukey whiskers; points represent individual biological replicates (n=3). Statistical

significance was determined using binomial generalized linear models with a logit link. **(F)** Relative community enrichment of all treatment groups (left), vehicle vs Mu-BiTE (middle) and vehicle vs combination (right) across communities. Bars indicate the proportion of cells contributed by each treatment group within a given community; values sum to 100% within each community. **(G)** Heatmap showing the proportion of CD8<sup>+</sup> T cells across cellular communities and treatment groups. **(H)** Relative community composition across tumor zones and treatment groups. Stacked area plots show the proportion of cellular communities within the Margin, Outer Core, Mid Core, and Deep Core zones, with each zone normalized to 100%. **(I-M)** Zone-specific distribution of selected cellular communities across treatment groups, weighted by the overall abundance of each community within the corresponding treatment group. (I) Tumor; (J) DC; (K) CAF-DC-CD8; (L) CAF; (M) DC-CAF. Lines represent the mean and shaded bands represent the SEM of biological triplicates (n=3).

Table S1: scFv sequences targeting AXL and PD-1

| Target | Ab name | scFv sequence (VL-(GGGS)4-VH-(GGGS)4) |
| --- | --- | --- |
| AXL | Enapotamab: scFv1 | EIVLTQSPGTLSLSPGERATLSCRASQSVSSSYLAWYQQKPGQAPRLLIYGASSRATGIPDRFSGSGSGTDFTLTISRLEPEDFAVYYCQQYGSSPYTFGQGTKLEIKGGGGSGGGGSGGGGSGGGGSEVQLLESGGGLVQPGGSLRLSCAASGFTFSSYAMNWVRQAPGKGLEWVSTTSGSGASTYYADSVKGRFTISRDNSKNTLYLQMNSLRAEDTAVYYCAKIWIAFDIWGQGTMVTVSSGGGGSGGGGSGGGGSGGGGS |
|  | YW327.6S2: scFv2 | DIQMTQSPSSLSASVGDRVTITCRASQDVSTAVAWYQQKPGKAPKLLIYSASFLYSGVPSRFSGSGSGTDFTLTISSLQPEDFATYYCQQSYTTPPTFGQGTKVEIKGGGGSGGGGSGGGGSGGGGSEVQLVESGGGLVQPGGSLRLSCAASGFSLSGSWIHWVRQAPGKGLEWVGWINPYRGYAYYADSVKGRFTISADTSKNTAYLQMNSLRAEDTAVYYCAREYSGWGGSSVGYAMDYWGQGTLVTVSSGGGGSGGGGSGGGGSGGGGS |
| PD-1 | Keytruda: scFv1 | EIVLTQSPATLSLSPGERATLSCRASKGVSTSGYSYLHWYQQKPGQAPRLLIYLASYLES GVPARFSGSGSGTDFTLTISSLEPEDFAVYYCQHSRDLPLTFGGGTKVEIKGGGGSGGGGSGGGGSGGGGSQVQLVQSGVEVKKPGASVKVSCKASGYTFTNYYMYWVRQAPGQGLEWMGGINPSNGGTNFNEKFKNRVTLTTDSSTTTAYMELKSLQFDDTAVYYCARRDYRFDMGFDYWGQGTTVTVSSGGGGSGGGGSGGGGSGGGGS |
|  | Nivolumab: scFv2 | EIVLTQSPATLSLSPGERATLSCRASQSVSSYLAWYQQKPGQAPRLLIYDASNRATGIPARFSGSGSGTDFTLTISSLEPEDFAVYYCQQSSNWPRTFGQGTKVEIKGGGGSGGGGSGGGGSGGGGSQVQLVESGGGVVQPGRSLRLDCKASGITFSNSGMHWVRQAPGKGLEWVAVIWYDGSKRYYADSVKGRFTISRDNSKNTLFLQMNSLRAEDTAVYYCATNDDYWGQGTLVTVSSGGGGSGGGGSGGGGSGGGGS |
| Murine PD-1 | Ab E: scFv1 | EIVLTQSPATLSLSPGERATLSCRASQSVSSSYLAWYQQKPGQAPRLLIYDASKRATGIPARFSGSGSGTDFTLTISLQPEDFAVYYCDQRNNWPLTFGGGQTKVEIKGGGGSGGGGSGGGSGGGGSGVQLVQSGAEVKKPGASVKVSKASGYTFTSNYMHVVRQAPGQGLEWMGIINPSEGSTGYAQKFQGRVTMTRDTSTSTVYMELSSLRSED TAVYYCAKEGVADGYGLVDVWGQGTMTVTVSSGGGGSGGGGSGGGGSGGGGS |
|  | Ab C: scFv2 | EIVLTQSPATLSLSPGERATLSCRASQSVSSSYLAWYQQKPGQAPRLLIYDASKRATGIPARFSGSGSGTDFTLTISLQPEDFAVYYCDQRNNWPLTFGGGQTKVEIKGGGGSGGGGSGGGSGGGGSGVQLVQSGAEVKKPGASVKVSKASGYTFTSNYMHVVRQAPGQGLEWMGIINPSEGSTGYAQKFQGRVTMTRDTSTSTVYMELSSLRSED TAVYYCAKEGVADGYGLVDVWGQGTMTVTVSSGGGGSGGGGSGGGGSGGGGS |

Table S2: Marker definitions used for cell phenotyping in murine multiplex imaging experiments

| Cell type | Defining markers |
| --- | --- |
| CD8 T cells | CD45<sup>+</sup> CD3<sup>+</sup> CD8<sup>+</sup> |
| CD4 T cells | CD45<sup>+</sup> CD3<sup>+</sup> CD4<sup>+</sup> |
| T Lymphocytes (unsorted) | CD45<sup>+</sup> CD3<sup>+</sup> |
| Regulatory T cells | CD45<sup>+</sup> CD3<sup>+</sup> CD4<sup>+</sup> FOXP3<sup>+</sup> |
| Activated T cells | Granzyme B<sup>+</sup> |
| Exhausted T cells | TOX<sup>+</sup> / LAG-3<sup>+</sup> / TIM-3<sup>+</sup> (any positive) |
| B cells | CD45<sup>+</sup> B220<sup>+</sup> |
| Neutrophils | CD45<sup>+</sup> Ly6G<sup>+</sup> |
| NK cells | CD45<sup>+</sup> T-bet<sup>+</sup> CD161c<sup>+</sup> |
| DCs | CD45<sup>+</sup> CD11c<sup>+</sup> |
| Myeloid cells | CD45<sup>+</sup> CD11b<sup>+</sup> |
| CAFs | αSMA<sup>+</sup> |
| Endotheliocytes | CD31<sup>+</sup> |
| Proliferating cells | Ki-67<sup>+</sup> |

Table S3: Primer sequences

| Primer name | Internal identifier | Sequence |
| --- | --- | --- |
| Primer A | F pFuse seq | GTTTTCTGTTCTGCGCCGTTACAG |
| Primer B | R pFuse seq | CCTCCAAATCAAGCCTCTACTTGAATCCT |
| Primer C | αAxl 1+2<br>gBlock R | GGTAGTAGATCTGTGGAGTGCCCAC |
| Primer D | F open AXL<br>pFuse | ATGGATGACATCCAAATGACTCAAAGCC |
| Primer E | R plas ran amp | CACTTGTCACGAATTCGATATCGGCC |
| Primer F | PD-1 amp F | TGTCACGAATTCGATATCGGCCATGGATGAAATTGTCCTCACCCAAAGCCCTG |
| Primer G | PD-1 amp R | GGAGGAGGAGGATCCGGTGGTGGAGGTAGTGACATCCAAATGACTCAAAGC |
| Primer H | F open PD-1<br>pFuse | ATGGATGAAATTGTCCTCACCCAAAGC |
| Primer I | AXL amp F | TCACGAATTCGATATCGGCCATGGATGACATCCAAATGACTCAAAGCCCATCCTCTCTCTCTG |
| Primer J | AXL amp R | GGAGGAGGAGGATCCGGTGGTGGAGGTAGTGAAATTGTCCTCACCCAAAG |
| Primer K | F open FC | CATCACCATCACCATCACTGAGTGCTAG |
| Primer L | R open FC | GCCTCTCCCTGTCTCCGGGTAAA |
| Primer M | F linker + AXL<br>amp | CTCTCCCTGTCTCCGGGTAAAGGTGGTGGAGGTTCGGGAGGAGGTG |
| Primer N | R linker + AXL<br>amp | GTTACGCAATGGATTACTGGGGCCAAGGTACTTTAGTTCATCACCATCACCATCACTGAGTGCT<br>AGCTGGC |
| Primer O | Plasmid amp F | AGATCTGTGGAGTGCCCACCTTG |
| Primer P | Plasmid amp R | GTCACGAATTCGATATCGGCCATGG |

Table S4: Mouse antibody panel

| Antibody | Clone | Channel | Dilution |
| --- | --- | --- | --- |
| CD45 | D3F8Q | 550 | 1:200 |
| CD3 | CAL57 | 550 | 1:200 |
| CD8 | D4W2Z | 550 | 1:50 |
| CD4 | EPR19514 | 647 | 1:50 |
| FOXP3 | FJK-16s | 647 | 1:50 |
| Granzyme B | EPR22645-206 | 550 | 1:1200 |
| TOX | E6I3Q | 550 | 1:200 |
| LAG-3 | CAL77 | 647 | 1:50 |
| TIM-3 | EPR22241 | 647 | 1:50 |
| B220 | RA3-6B2 | 550 | 1:100 |
| Ly6G | E6Z1T | 550 | 1:100 |
| T-bet | E4I2K | 647 | 1:50 |
| CD161c | E6Y9G | 550 | 1:100 |
| CD11c | D1V9Y | 647 | 1:80 |
| CD11b | EPR1344 | 750 | 1:250 |
| αSMA | SP171 | 750 | 1:500 |
| CD31 | RM1006 | 550 | 1:1000 |
| Ki-67 | B56 | 550 | 1:600 |
